# A heteromeric nicotinic acetylcholine receptor promotes sleep by relaying GABAergic signals within a locus of motor and sensory integration

**DOI:** 10.1101/2024.10.02.616374

**Authors:** Tomohiro Yumita, Hassan Ahamed, Hayden A. M. Hatch, Ian Cossentino, Charalambia Louka, Nicholas Stavropoulos

## Abstract

Locomotor inactivity and reduced sensory responsiveness are defining attributes of sleep, yet the underlying mechanisms are not well understood. In particular, the molecular and circuit mechanisms by which sleep-regulatory signals from the brain restrict movement and sensation remain largely ill-defined. Here we identify a nicotinic acetylcholine receptor (nAChR) that promotes sleep in *Drosophila* through its expression in GABAergic neurons of the ventral nerve cord (VNC), a center for motor and sensory systems. Biochemical, genetic, and pharmacological manipulations indicate that a heteromeric nAChR containing the α1 and β1 subunits promotes sleep by coupling cholinergic input to GABA release in the VNC and the likely inhibition of motor neurons, sensory afferents, or both. The functional parallels of the VNC and the mammalian spinal cord suggest that disruptions of analogous inhibitory circuits in humans may impair suppression of behavioral activity and sensory inputs during sleep and contribute to sleep disorders.

## INTRODUCTION

Sleep is a restorative state critical for normal physiology and brain function and is highly conserved across the animal kingdom.^1,2^ Locomotor inactivity and reduced responsiveness to sensory stimuli are universal attributes of sleep, yet their molecular and circuit basis is not well understood. While molecules and neurons that regulate sleep have been intensely investigated within the brains of various animal models including *Drosophila*,^3–6^ the precise mechanisms by which sleep-regulatory signals suppress motor and sensory systems—which include neurons and projections residing outside of the brain—are less well defined.

Manipulations of sensory neurons and muscle can strongly influence sleep duration,^7–11^ suggesting that regulation of motor and sensory pathways is critical for sleep-wake cycles. Faithful communication between the brain and sensory and motor systems outside of the brain is essential for the proper regulation of sleep, as indicated by spinal cord lesions that perturb sleep.^12,13^ The neuronal and molecular mechanisms by which sleep-regulatory signals impinge on motor and sensory targets are clinically significant. Deficits in suppressing motor activity and sensory responsiveness are associated with pathological disruptions in sleep, including in REM sleep behavior disorder, various parasomnias, neurodegenerative conditions, and neurodevelopmental disorders such as autism.^14–16^

Acetylcholine and its receptors have critical and conserved functions in regulating sleep-wake cycles.^17^ Acetylcholine was among the first neurotransmitters linked to sleep regulation,^18^ and numerous studies have defined cholinergic nuclei and circuits important for sleep and arousal in both vertebrates and invertebrates.^4,5,19,20^ In humans, mutations in several subunits of the nicotinic acetylcholine receptor (nAChR) are linked to sleep-related hypermotor epilepsy,^21–23^ a disorder characterized by ectopic motor activity, dystonic episodes, and ambulatory wandering during sleep.^24^ A major barrier to understanding the impact of cholinergic neurotransmission on sleep is the complexity of acetylcholine receptors. In particular, nAChRs are formed by the pentameric assembly of various subunits expressed in overlapping distributions within the nervous system,^25^ posing a formidable challenge to understanding which discrete nAChRs assemble in particular neuronal populations to influence sleep-wake behavior. In *Drosophila*, the repertoire of nAChR subunits comprises seven ligand-binding α subunits (α1-α7) and three structural β subunits (β1-β3).^26–35^ Prior studies in the fly have shown that individual nAChR subunits can regulate sleep positively and negatively,^36–40^ but the identity of nAChR heteromers that influence sleep and their neuronal sites of action are largely unknown.

*γ*-aminobutyric acid (GABA) and its receptors have a similarly prominent role in regulating sleep.^4,5,19,20,41^ GABAA receptor agonists are the predominant class of sleep-promoting drugs in clinical use and have strong hypnotic effects in both mammals and flies.^42–44^ Mutations that impair GABA catabolism and increase GABA levels promote the consolidation and duration of sleep,^45^ and several populations of GABAergic neurons that regulate sleep have been described in the *Drosophila* brain.^46–51^ The potent hypnotic effects of GABA agonists and the broad expression of GABA receptors throughout the nervous system, including in neurons outside of the brain,^52–55^ suggest that additional, uncharacterized populations of GABAergic neurons contribute to the regulation of sleep-wake cycles.

Here, we identify a sleep-promoting function for a heteromeric nicotinic acetylcholine receptor, containing the α1 and β1 subunits, that acts in GABAergic neurons of the ventral nerve cord (VNC), a structure that governs motor output and relays motor and sensory signals to and from the brain. We propose that α1/β1 nAChRs expressed in GABAergic VNC neurons couple excitatory cholinergic input from the brain to the inhibition of targets in the VNC, including motor and sensory systems. Our findings implicate a mechanism in which sleep signals from central circuits activate inhibitory neurons outside of the brain to promote and maintain the sleep state by restricting behavioral activity, suppressing sensory inputs, or through both mechanisms.

## RESULTS

### The α1 and β1 subunits of the nicotinic acetylcholine receptor promote the duration and consolidation of sleep

To identify neurotransmitter receptors that regulate sleep, we used the Gal4/UAS system^56^ to direct RNAi against several metabotropic receptors and the complete repertoire of pentameric ligand-gated ion channels, including nicotinic acetylcholine receptors (nAChRs), *γ*-aminobutyric acid A (GABAA) receptors, and histamine- and glutamate-gated chloride channels (Figure 1A). We depleted these receptors with the pan-neuronal *elav^c^*^155^–*Gal4* driver^57^ and 70 RNAi lines targeting 32 genes. Neuronal expression of three RNAi lines sharply reduced sleep duration, to less than two standard deviations from the mean of all screened lines (Figure 1A). These RNAi lines targeted nAChR subunits α1, α4, and β1 (Figure 1A). A missense mutation within α4 was previously described to cause short sleep,^38^ and hence we focused further studies on α1 and β1.

**Figure 1.**
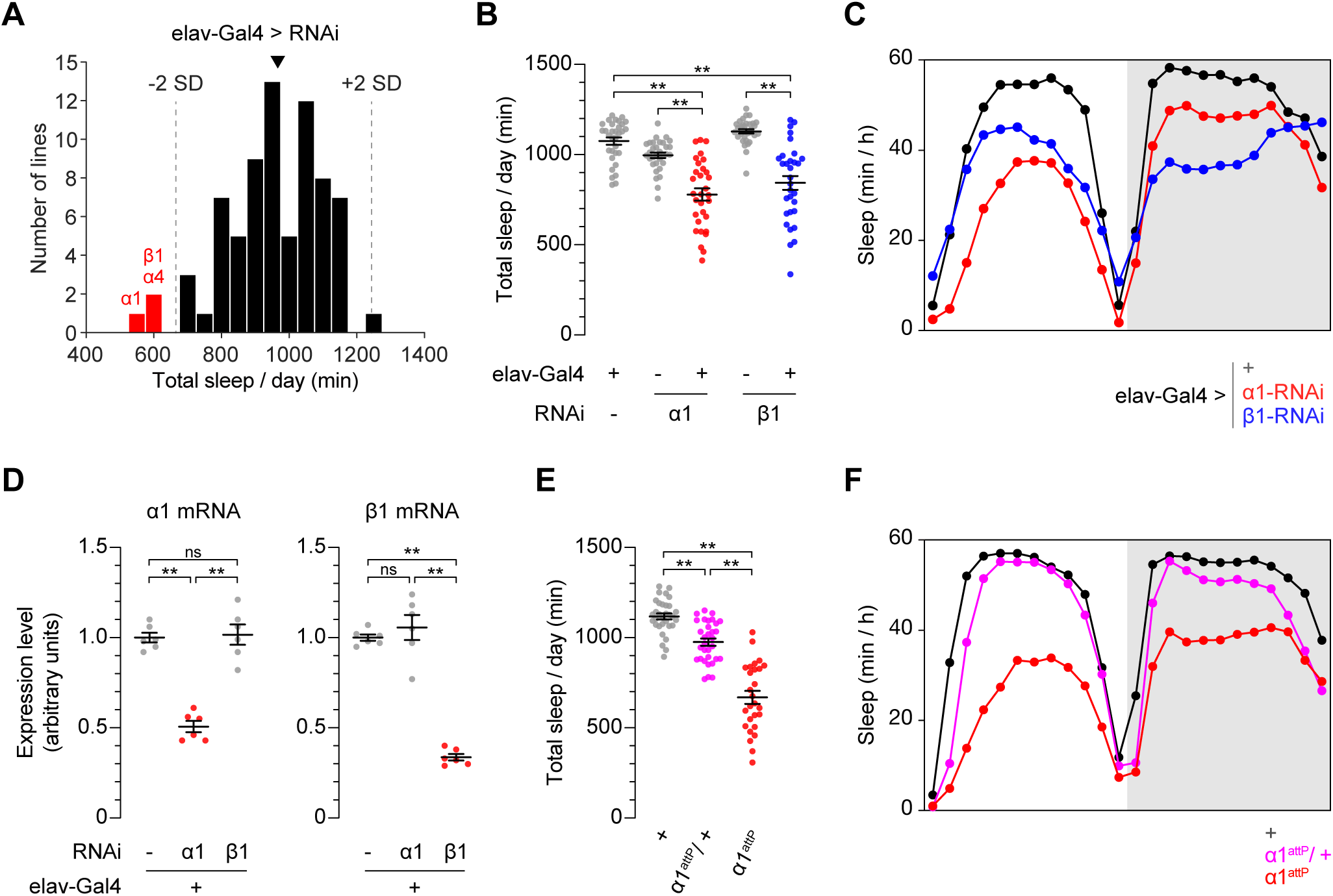
nAChR subunits α1 and β1 regulate sleep. **(A)** Histogram of total sleep for lines expressing pan-neuronal RNAi against neurotransmitter receptors. Arrowhead indicates average total sleep of *elav^C^*^155^–*Gal4; UAS-*dcr2 control animals lacking RNAi. Dotted lines indicate two standard deviations from the mean of all screened lines. **(B)** Pan-neuronal α1 or β1 RNAi elicits short sleep. n = 31-32. **(C)** Average sleep profiles of animals shown in (B). **(D)** Quantification of α1 or β1 transcripts in adult heads by qRT-PCR. **(E)** Homozygous *α1^attP^* and heterozygous *α1^attP^/+* animals exhibit short sleep. n = 27-32. **(F)** Average sleep profiles of animals shown in (E). **(B, D and E)** Bars indicate mean ± SEM. One-way ANOVA and Tukey post hoc tests; **p < 0.01; ns, not significant.

We confirmed and characterized the sleep phenotypes arising from α1 and β1 depletion after backcrossing RNAi transgenes. Pan-neuronal RNAi against α1 or β1 strongly curtailed total sleep duration with respect to controls bearing only Gal4 or RNAi transgenes (Figures 1B and 1C). Two additional RNAi lines targeting a non-overlapping region of the α1 transcript similarly shortened sleep when expressed in neurons (Figures S1A and S1B), suggesting that these short sleep phenotypes reflected α1 depletion rather than off-target effects. For both α1 and β1, neuronal RNAi reduced sleep during both daytime and nighttime and did not alter its circadian oscillation in light-dark cycles (Figures 1C and S1B–S1D). Short sleep caused by α1 or β1 RNAi was fragmented, as indicated by decreased sleep bout length and an increase in the number of daily sleep bouts (Figures S1E and S1F). Neuronal α1 RNAi did not significantly alter locomotor activity during wakefulness, indicating that reduced sleep was independent of locomotor activity (Figure S1G). While neuronal RNAi against β1 increased locomotor activity in a fraction of animals, these changes were not correlated with reduced sleep (Figures S1G and S1H).

To assess the efficacy and specificity of α1 and β1 RNAi, we used qRT-PCR to measure α1 and β1 mRNA levels in heads of animals expressing neuronal α1 or β1 RNAi. Compared to controls bearing only *elav-Gal4*, neuronal α1 RNAi decreased α1 mRNA levels by 49%, and β1 RNAi reduced β1 mRNA levels by 66% (Figure 1D). α1 transcript levels were not altered by β1 RNAi and vice versa, indicating that depletion of each transcript was specific and did not elicit compensatory upregulation of the other.

To independently assess whether reduced activity of α1 and β1 impacts sleep, we generated mutant alleles for both genes with the CRISPR/Cas9 system.^58^ *α1^attP^* and *β1^attP^* bear deletions of first coding exons and insertions of attP sites, yielding presumed null alleles that enable subsequent genetic manipulations of these loci (Figure S2A). *α1^attP^* animals were viable and exhibited severely curtailed sleep (Figures 1E and 1F), recapitulating the phenotype caused by neuronal α1 RNAi. Sleep was also reduced in *α1^attP^/+* heterozygotes (Figures 1E and 1F), consistent with a dose-dependent influence on sleep and the effects of neuronal α1 depletion. Like animals expressing neuronal α1 RNAi, *α1^attP^* and *α1^attP^/+* animals exhibited short and fragmented sleep in daytime and nighttime (Figures S2B–S2E) and were not hyperactive (Figure S2F). *β1^attP^* animals were short-lived and rarely survived the duration of the sleep assay, precluding assessment of their sleep phenotypes. The phenotypes of *α1^attP^* and *α1^attP^/+* animals, together with those of animals expressing neuronal α1 or β1 RNAi, indicate that α1 and β1 are required in neurons for normal sleep duration and consolidation.

The short sleep phenotypes caused by loss-of-function manipulations of α1 and β1 were surprising, as nAChRs are excitatory and broadly expressed throughout the nervous system.^59–61^ Reduced α1 or β1 activity might thus be expected to globally dampen neuronal excitation and mimic the effects of hypnotic drugs such as gaboxadol, which augment neuronal inhibition and promote sleep.^42–44^ Further investigation, as described below, suggests that α1 and β1 act in inhibitory GABA-releasing neurons to promote sleep, suggesting that their reduced activity causes neuronal disinhibition, offering a resolution to this paradox.

### α1 and β1 impact baseline sleep regulation

Sleep is regulated by circadian and homeostatic mechanisms.^62^ In *Drosophila*, cholinergic neurotransmission is implicated in both circadian rhythms and the homeostatic regulation of sleep. Circadian pacemaker neurons receive cholinergic input and express nAChRs^63–66^, and broad cholinergic activation triggers sleep loss and subsequent homeostasis^9^. We first assessed whether neuronal depletion of α1 or β1 alters circadian rhythmicity, by entraining animals in light-dark cycles and measuring locomotor activity subsequently in constant darkness. Animals expressing neuronal α1 or β1 RNAi exhibited behavioral periods indistinguishable from those of control animals, suggesting that circadian oscillations are intact, with weaker locomotor rhythms characteristic of short sleep mutants^67,68^ (Figure 2A).

**Figure 2:**
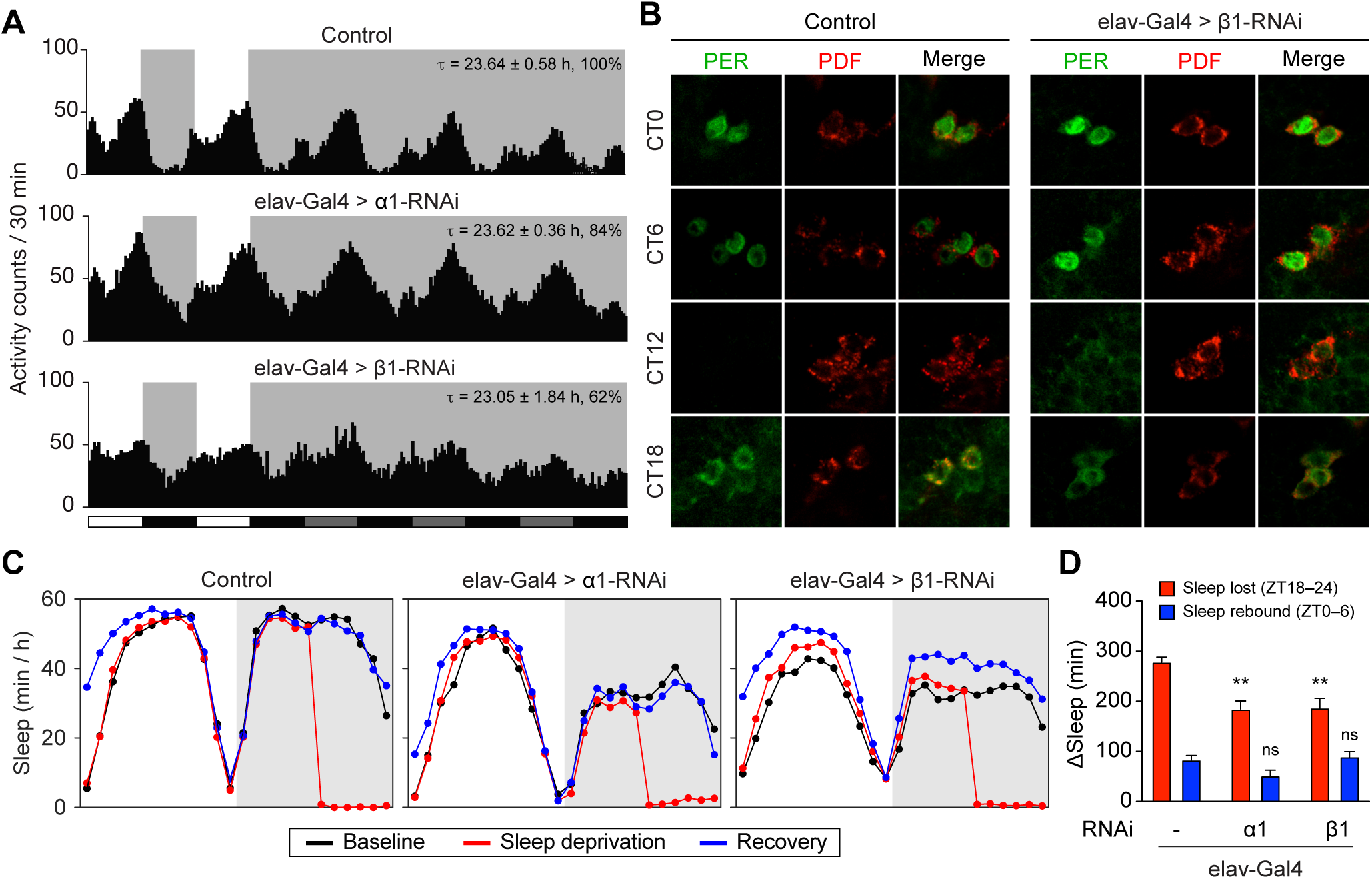
Depletion of α1 and β1 does not alter the circadian and homeostatic regulation of sleep. **(A)** Average locomotor activity of *elav^C^*^155^–*Gal4* control animals or animals expressing pan- neuronal RNAi against α1 or β1. Two days of light-dark cycles and three days of constant darkness are shown. Periodicity (*τ*) and percentage of rhythmic animals are indicated at upper right. n = 13-24. **(B)** Oscillations of Period (PER) abundance and localization in pigment- dispersing factor (PDF)-expressing small ventrolateral neurons (sLNvs) are indistinguishable between *elav^C^*^155^–*Gal4* controls and animals expressing pan-neuronal β1 RNAi. **(C)** Average sleep profiles for baseline (black), sleep deprivation (red), and recovery (blue) days for *elav^C^*^155^–*Gal4; UAS-dcr2/+* control animals or animals expressing pan-neuronal RNAi against α1 or β1. Note 6h of sleep deprivation at ZT18–24 and rebound on the subsequent recovery day. **(D)** Quantification of sleep lost (ZT18–24) and sleep recovered (ZT0–6) for genotypes in **(C)**. n = 35-38. Mean ± SEM is shown. Sleep lost and rebound are compared to control. One-way ANOVA and Tukey post hoc tests. **p < 0.01; ns, not significant.

We further assessed whether neuronal β1 RNAi, which reduces the strength of locomotor rhythms more than α1 RNAi, alters the molecular oscillations of the circadian clock. We performed immunohistochemical staining for the core clock protein Period (Per), whose oscillations in circadian pacemaker neurons underlie rhythmic behavior.^69–72^ Control brains and brains expressing neuronal RNAi against β1 exhibited indistinguishable oscillations in Per localization and abundance in pigment-dispersing factor (PDF)-expressing small ventrolateral neurons (sLNvs), with low PER levels at CT12, accumulation of PER in the cytoplasm at CT18, strong PER signal in nuclei at CT0, and PER degradation at CT6 through CT12 (Figure 2B). The molecular oscillations of the circadian clock are thus intact in animals with reduced β1 levels, paralleling normal behavioral oscillations of sleep-wake cycles (Figure 2A). Together, these data suggest that neuronal α1 and β1 depletion does not alter the circadian regulation of sleep.

To determine whether α1 or β1 depletion alters sleep homeostasis, we next assessed the response to sleep deprivation in animals expressing neuronal α1 or β1 RNAi. Intermittent mechanical stimulation for six hours at the end of the night elicited near-complete sleep deprivation and increased sleep the following day in wild-type animals (Figures 2C and 2D). While animals expressing neuronal RNAi against α1 and β1 lost smaller amounts of sleep during sleep deprivation due to lower baseline sleep levels, they exhibited sleep rebound similar to that of control animals (Figures 2C and 2D), suggesting that sleep homeostasis is intact. These data indicate that α1 and β1 are important for the duration and maintenance of sleep, but act independently of its circadian and homeostatic regulation.

### α1 and β1 assemble into a heteromeric nAChR and act through a common population of neurons to impact sleep

nAChRs are pentameric ion channels that assemble from acetylcholine-binding α subunits and structural β subunits^73^. While the subunit composition of *Drosophila* nAChRs *in vivo* remains largely unknown, the similar sleep phenotypes caused by manipulations of α1 or β1 indicate that these subunits may assemble into a heteromeric receptor essential for sleep. While α1 and β1 are known to be expressed widely in the nervous system,^61^ their co-localization and physical interactions have not been assessed.

To bypass the lack of antibodies effective for co-staining of α1 and β1, we introduced epitope tags within the *α1* and *β1* loci. We used CRISPR/Cas9 targeting and homology directed to insert smGFP-V5 and smGFP-FLAG tags, each containing 10 epitope tags within a GFP scaffold^74^, into the large intracellular loops of α1 and β1, respectively (α1^V5^ and β1^FLAG^, Figures 3A and S3). Immunostaining of animals bearing α1^V5^ and β1^FLAG^ revealed that α1 and β1 are distributed broadly in synaptic neuropil and somata throughout the brain and VNC, in overlapping but not identical patterns (Figures 3B and 3C). α1 and β1 colocalized at synaptic puncta throughout the nervous system, including within the optic lobe, antennal lobe, subesophageal zone, and the VNC (Figure 3C). Co-staining with 4’,6-diamidino-2-phenylindole (DAPI) indicated that α1 and β1 signals were excluded from cell nuclei, as expected for these receptor proteins (Figure 3C). These results indicate that α1 and β1 colocalize broadly within the nervous system and at the level of synapses.

**Figure 3:**
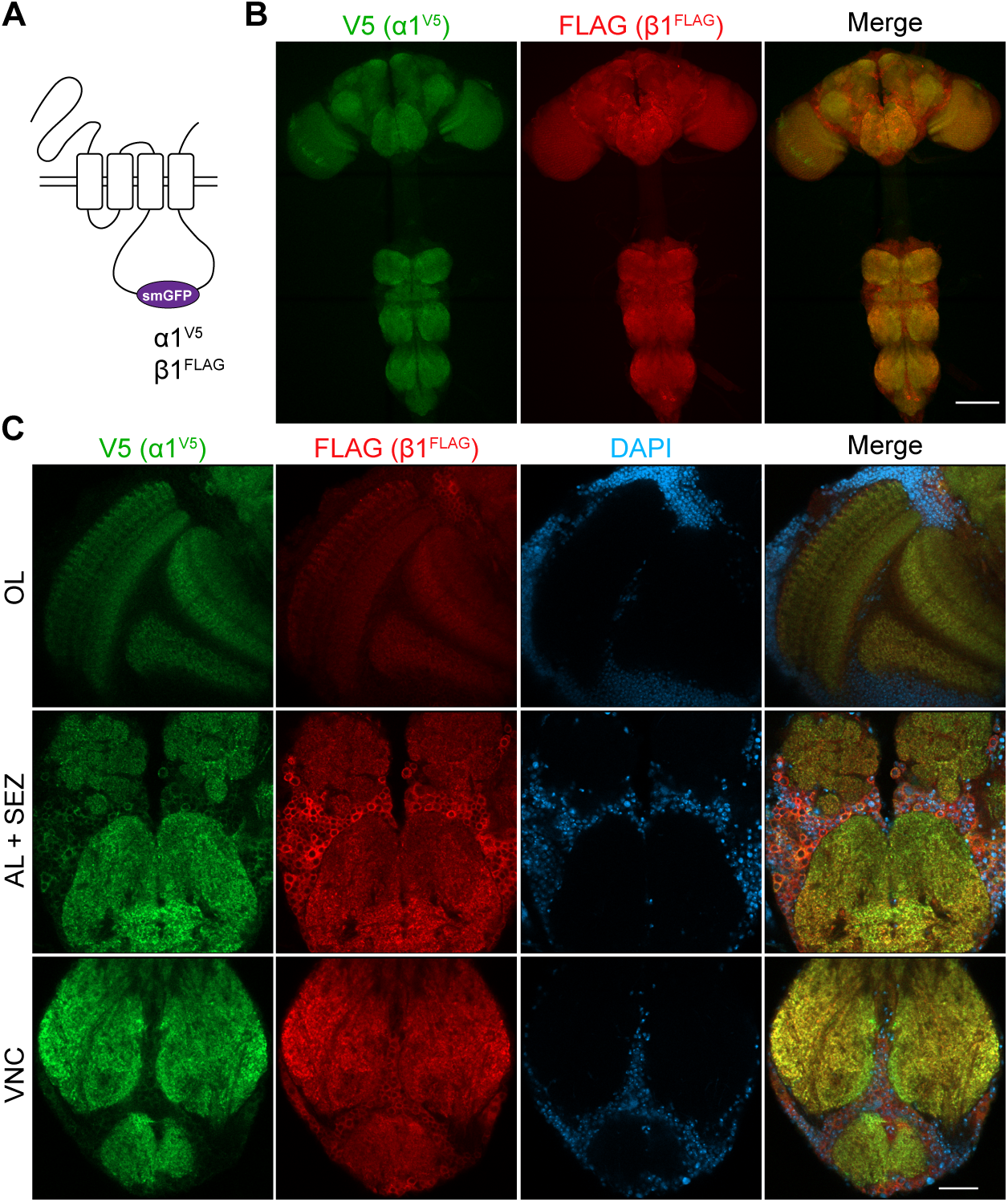
Endogenous tagging and localization of α1 and β1. **(A)** Strategy for endogenously tagging α1 and β1 with high affinity spaghetti-monster GFP (smGFP) tags inserted in the large intracellular loop between the third and fourth transmembrane domains. **(B)** Maximal projections of the brain and ventral nerve cord of a *β1^FLAG^ α1^V5^/+* animal. **(C)** Single confocal slices of the optic lobe (OL), antennal lobe (AL), subesophageal zone (SEZ), and ventral nerve cord (VNC) of a *β1^FLAG^ α1^V5^/+* animal. Note overlap of α1^V5^ and β1^FLAG^ signal in synaptic neuropil and somata and exclusion from nuclei. Scale bars represent 100 µm for **(B)** and 25 µm for **(C)**.

We next tested whether α1 and β1 assemble into heteromeric nAChRs *in vivo*, by performing reciprocal co-immunoprecipitations from animals bearing endogenously tagged α1^V5^ and β1^FLAG^. We found that immunoprecipitation of α1^V5^ from head lysates retrieved β1^FLAG^ and vice versa, indicating that α1 and β1 interact physically within the brain (Figure 4A). This result indicates that α1 and β1 assemble into heteromeric nAChRs *in vivo*.

**Figure 4:**
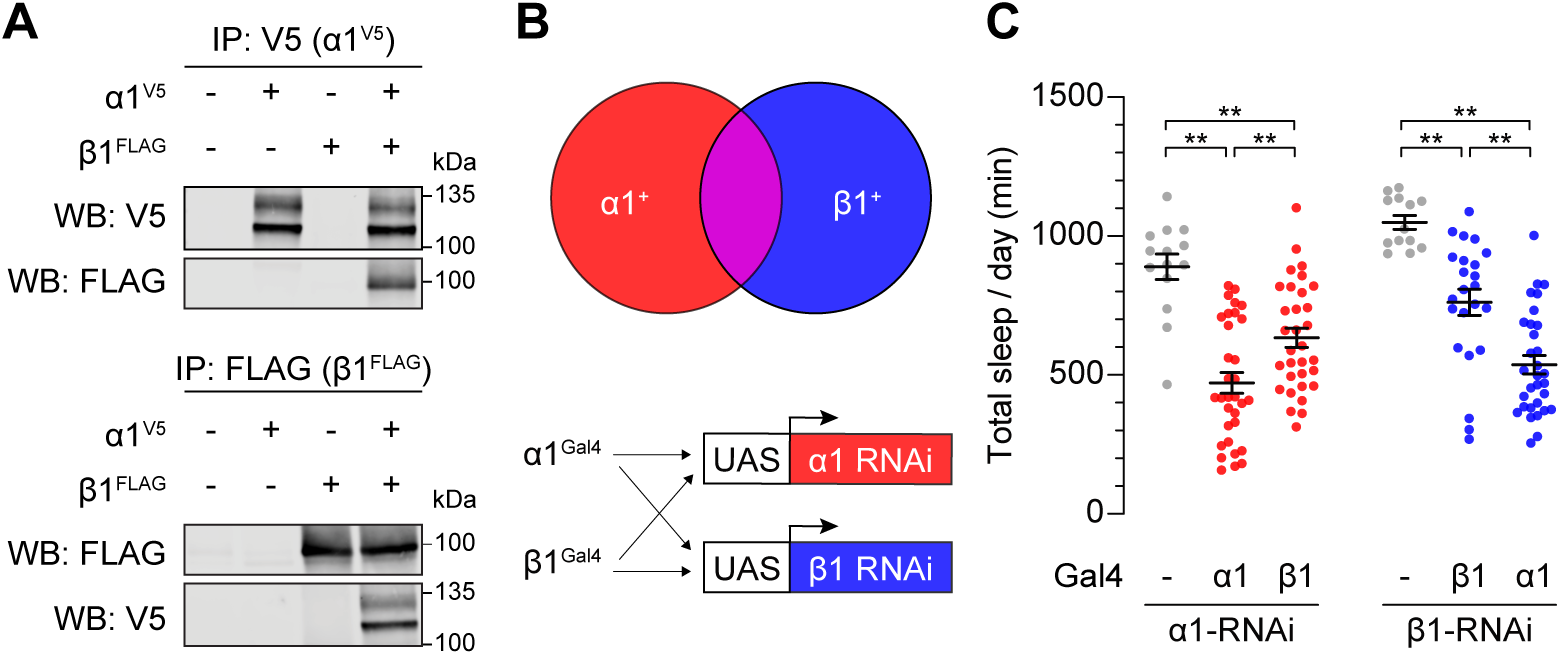
α1 and β1 form a functional heteromeric nAChR that regulates sleep. **(A)** α1 and β1 assemble into a heteromeric nAChR *in vivo*. Reciprocal co-immunoprecipitations (IP) and Western blots (WB) of α1 and β1 from head lysates prepared from *β1^FLAG^ α1^V5^* heterozygotes or controls of indicated genotypes. **(B)** RNAi strategy for targeting neurons that express α1, β1, or both. **(C)** α1 and β1 regulate sleep through neurons in which they are co- expressed. n = 13-32. Bars indicate mean ± SEM. One-way ANOVA and Tukey post hoc tests; **p < 0.01.

To determine whether heteromeric nAChRs containing α1 and β1 promote sleep, we sought to obtain genetic access to neurons that natively express these receptor subunits. We first assessed Gal4 lines from the FlyLight collection^75^ that bear upstream and intronic fragments from the *α1* and *β1* loci. We evaluated six Gal4 lines for *α1* and two for *β1*, using them to drive α1 or β1 RNAi and assess the consequences on sleep. None of these manipulations altered sleep (Figure S4A), suggesting that these Gal4 lines do not recapitulate α1 and β1 expression. To obtain Gal4 drivers that faithfully recapitulate α1 and β1 expression, we introduced Gal4 into the *α1* and *β1* loci. We generated *α1^Gal4^* by inserting a T2A-Gal4 cassette into the third intron of *α1* using the MiMIC/Trojan system,^76^ a manipulation predicted to yield Gal4 expression from the *α1* locus and to truncate the α1 protein to a nonfunctional form (Figure S4B). To generate *β1^Gal4^*, we integrated a Gal4-bearing plasmid into the *attP* site of *β1^attP^* with the *Φ*C31 system^77^ and subsequently excised the 3*×*P3-GFP selection marker, yielding Gal4 fused to the endogenous β1 start codon and a predicted *β1* null allele (Figure S4C).

To assess whether *α1^Gal4^* and *β1^Gal4^* recapitulate expression in neurons through which α1 and β1 influence sleep, we measured sleep in animals in which *α1^Gal4^* and *β1^Gal4^* were used to respectively deplete α1 and β1 (Figure 4B). Expressing α1 RNAi with *α1^Gal4^* elicited short sleep comparable to that caused by pan-neuronal α1 depletion (Figure 4C). Similarly, expressing β1 RNAi with *β1^Gal4^* recapitulated the short sleep phenotype of pan-neuronal β1 RNAi (Figure 4C). Thus, *α1^Gal4^* and *β1^Gal4^* functionally recapitulate α1 and β1 expression in neurons through which these proteins impact sleep.

The similar short-sleeping phenotypes caused by pan-neuronal depletion of α1 or β1, together with their assembly into a heteromeric nAChR *in vivo*, prompted us to test whether α1 and β1 impact sleep through an overlapping population of neurons (Figure 4B). Consistent with this possibility, using *β1^Gal4^* to drive α1 RNAi elicited short sleep similar to that caused by depleting α1 pan-neuronally or in α1-expressing neurons (Figure 4C). Conversely, using *α1^Gal4^* to deplete β1 specifically in neurons that express α1 elicited short sleep comparable to that caused by β1 depletion in all neurons or in β1-expressing neurons (Figure 4C). These reciprocal genetic manipulations indicate that α1 and β1 regulate sleep through a population of neurons in which they are co-expressed. Together with the overlapping distribution of α1 and β1 in the nervous system and their physical associations *in vivo* (Figures 3B, 3C, and 4A), these data strongly suggest that α1 and β1 assemble into a heteromeric nAChR essential for sleep regulation.

### α1 and β1 promote sleep through their expression in GABAergic neurons

To further define neuronal populations through which α1 and β1 regulate sleep, we performed screens in which we expressed α1 or β1 RNAi in various neuronal populations using the Gal4/UAS system. We screened 360 Gal4 lines with α1 RNAi and 76 Gal4 lines with β1 RNAi and found that two *vGat-Gal4* transgenes caused the most severe short-sleeping phenotypes (Figures 5A, 5B, and 7A). *vGat* encodes the sole vesicular GABA transporter in *Drosophila* and marks GABAergic neurons,^78^ suggesting that α1 and β1 function together in GABAergic neurons to promote sleep. Given that increased GABAergic tone promotes sleep,^42–44^ reduced excitatory input to sleep-regulatory GABAergic neurons caused by depletion of α1/β1-containing nAChRs might be expected to prolong wakefulness by reducing GABA release, offering a potential explanation for the counterintuitive short sleep phenotypes caused by α1 and β1 depletion.

**Figure 5:**
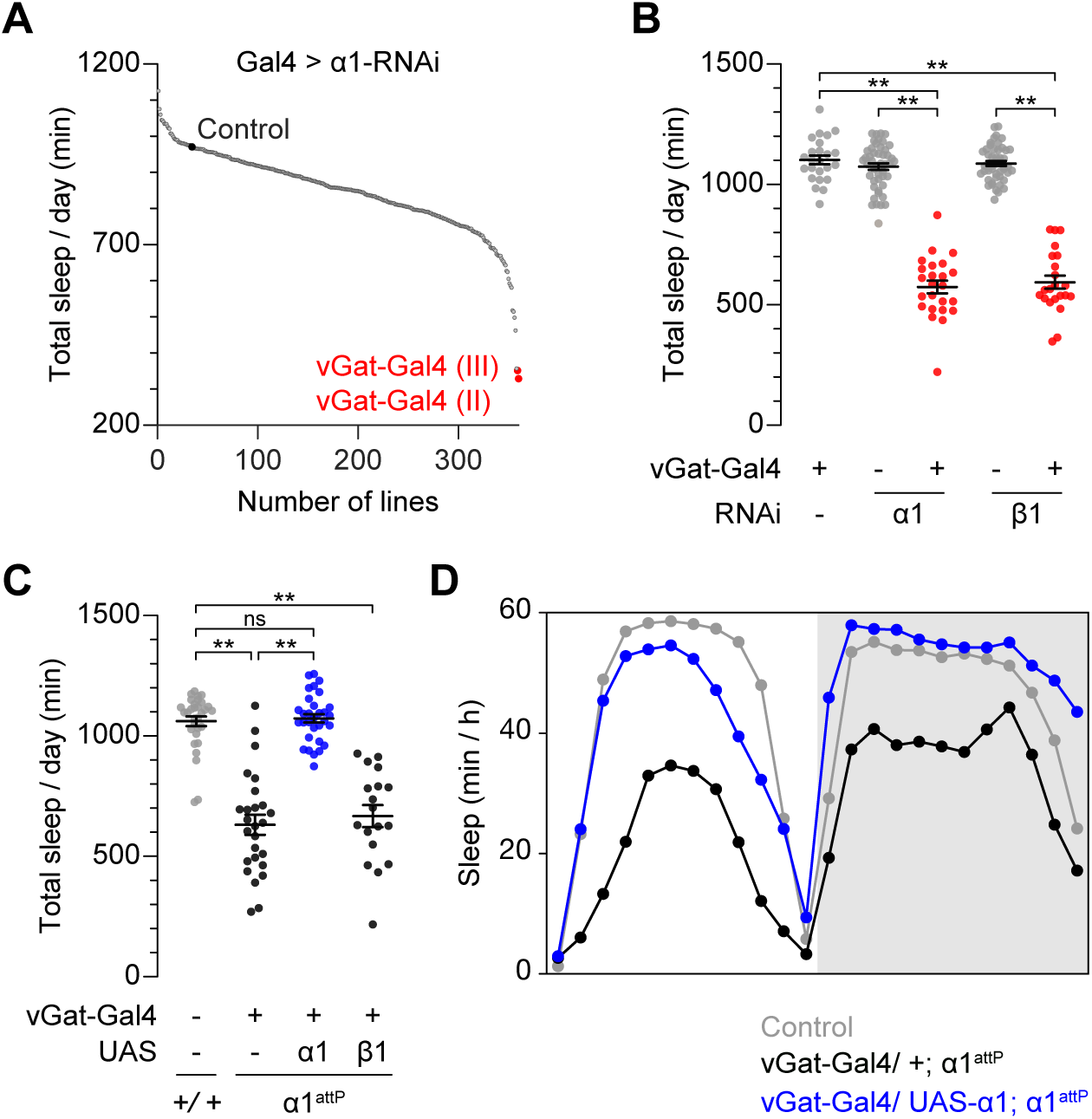
α1 and β1 regulate sleep through GABAergic neurons. **(A)** Total sleep for Gal4 lines expressing α1 RNAi. Black dot, *UAS-α1-RNAi; UAS-dcr2* control animals lacking Gal4. *vGat-Gal4 (II)* and *vGat-Gal4 (III)* are 2nd and 3rd chromosome insertions respectively. **(B)** Depletion of α1 or β1 in neurons expressing *vGat-Gal4* elicits short sleep. n = 22-48. **(C)** α1 expression with *vGat-Gal4* restores sleep to homozygous *α1^attP^* mutants. Animals are heterozygous for *vGat-Gal4*, *UAS-α1*, and/or *UAS-β1* transgenes as indicated. n = 18-32. **(D)** Average sleep profiles of animals shown in **(C)**. **(B and C)** Bars indicate mean ± SEM. One-way ANOVA and Tukey post hoc tests; **p < 0.01; ns, not significant.

To test whether α1 and β1 expression in GABAergic neurons is sufficient to account for their impact on sleep, we first assessed whether expressing of α1 with *vGat-Gal4* could restore sleep to short-sleeping *α1^attP^* mutants. While homozygous *α1^attP^* animals bearing *vGat-Gal4* alone exhibited severe short-sleeping phenotypes, *α1^attP^* animals bearing both *vGat-Gal4* and *UAS-α1* slept indistinguishably from wild-type controls (Figures 5C and 5D), indicating that α1 expression with *vGat-Gal4* was sufficient to restore normal sleep. In contrast, β1 expression using *vGat-Gal4* failed to restore sleep to *α1^attP^* animals (Figure 5C), indicating that complementation by α1 was specific. Together, these results strongly suggest that α1 expression in GABAergic neurons is both necessary and sufficient to account for the impact of α1 on sleep. While β1 expression using *vGat-Gal4* restored viability to homozygous *β1^attP^* animals, short sleep persisted in these animals, suggesting that β1 has essential functions in additional sleep-regulatory neurons that do not express *vGat-Gal4*, or that manipulations with *vGat-Gal4* do not recapitulate β1 expression in GABAergic neurons.

### The GABAA receptor agonist gaboxadol rescues the short sleep caused by α1 and β1 depletion

Functional manipulations of α1 and β1 with *vGat-Gal4* suggest that excitatory input through heteromeric α1/β1-containing nAChRs in GABAergic neurons promotes sleep, and that decreased cholinergic input to these neurons causes short sleep by reducing GABA release. To test this hypothesis, we examined whether augmenting GABAA receptor activity pharmacologically could restore sleep to animals in which α1 or β1 was depleted. We administered the GABAA receptor agonist 4,5,6,7-tetrahydroisoxazolopyridin-3-ol (gaboxadol) through feeding and assessed the effects on sleep. We used low gaboxadol concentrations that have negligible effects on sleep in wild-type animals (0.005 mg/ml and 0.01 mg/ml),^42^ reasoning that animals deficient for GABAergic signaling in sleep-regulatory neurons would be highly sensitive to gaboxadol. As expected, control animals fed 0.01 mg/ml or 0.005 mg/ml gaboxadol exhibited little or no increase in sleep, respectively (Figures 6A and 6B and S5A). In contrast, the same concentrations of gaboxadol restored daytime sleep to near wild-type levels in animals expressing α1 or β1 RNAi under the control of *vGat-Gal4* (Figures 6A and 6B); nighttime sleep was not measurably increased by gaboxadol (Fig S5B). Notably, these animals were most sensitive to the lowest concentration of gaboxadol (0.005 mg/ml), as indicated by sharply increased sleep with respect to vehicle controls. Locomotor activity during wakefulness was unchanged in animals exposed to gaboxadol, indicating that increased sleep did not reflect locomotor deficits (Figure 6C). Consistent with these findings, the same concentrations of gaboxadol strongly increased daytime sleep duration in animals expressing pan-neuronal α1 or β RNAi but did not affect their locomotion (Figure S6A–E). Collectively, these results indicate that increased GABAA receptor activity can bypass the effects of depleting α1 and β1, consistent with a critical function of heteromeric α1/β1-containing nAChRs in eliciting GABA release from sleep-regulatory GABAergic neurons.

**Figure 6:**
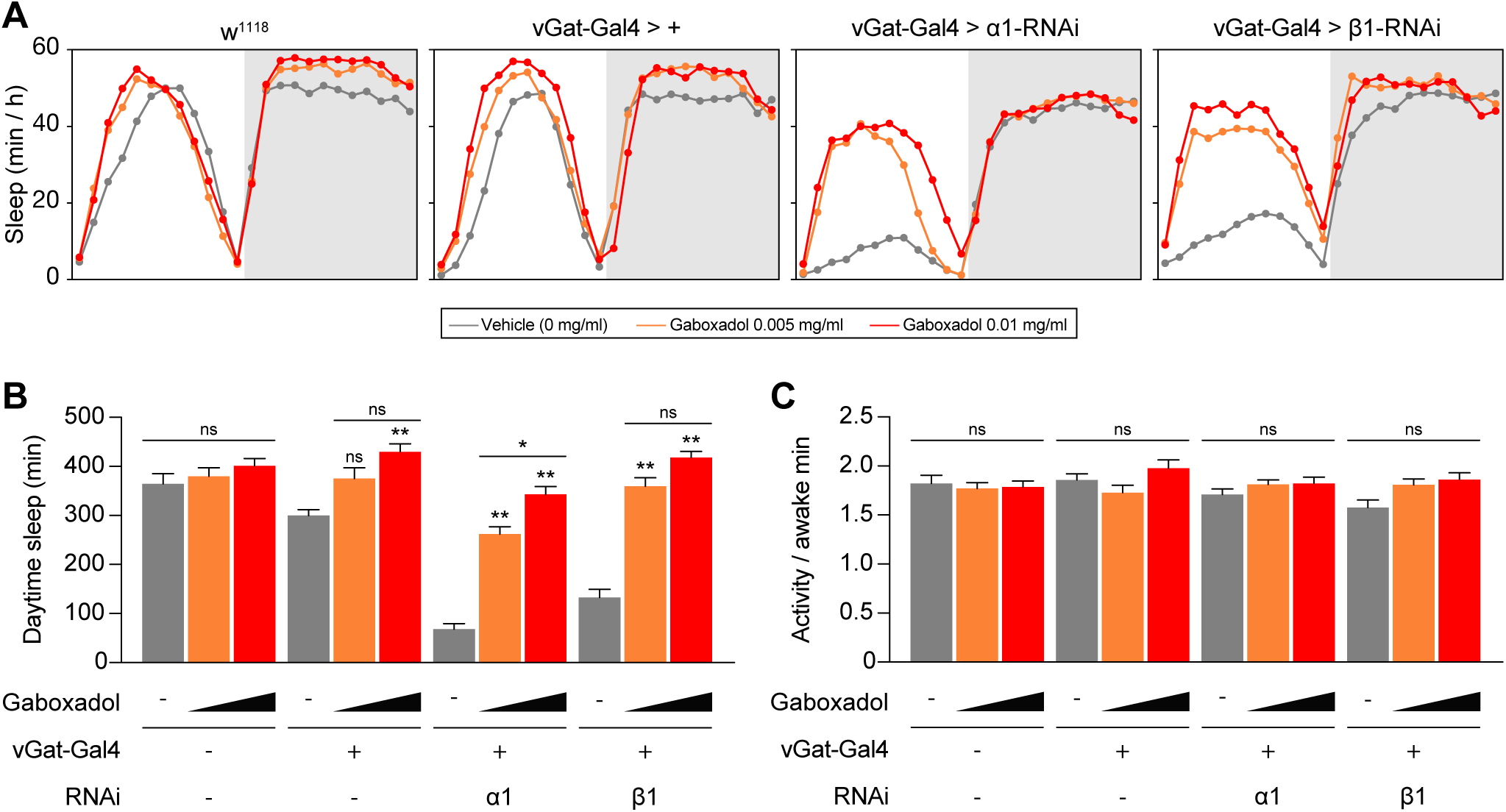
The GABAA receptor agonist gaboxadol complements the sleep deficits of animals expressing α1 and β1 RNAi under *vGat-Gal4* control. **(A)** Average sleep profiles of gaboxadol-treated animals of indicated genotypes. **(B)** Short sleep caused by α1 or β1 depletion using *vGat-Gal4* is rescued by low concentrations of gaboxadol. **(C)** Locomotor activity is not altered by gaboxadol treatment. For **(B and C)**, mean ± SEM is shown. n = 21-22. Two-way ANOVA and Tukey post hoc tests; *p < 0.05; **p < 0.01; ns, not significant.

### α1 and β1 function in GABAergic neurons in the ventral nerve cord

Many neurons and genes previously found to impact sleep in *Drosophila* reside or act in the brain.^4,19^ Intriguingly, in our neuroanatomical screen using various Gal4 drivers to express β1 RNAi, we found that a Gal4 insertion within the *teashirt* (*tsh*) locus (*tsh-Gal4*) strongly curtailed sleep when used to drive β1 RNAi (Figure 7A). *tsh* expression in the nervous system is restricted mainly to the VNC^79,80^ (Figure 7D), suggesting that the effects of β1 RNAi on sleep might arise outside the brain. We backcrossed *tsh-Gal4* to allow careful behavioral comparisons and tested whether sleep was altered by depleting α1 and β1 in VNC neurons using *tsh-Gal4*. Pupal lethality was observed in animals bearing *tsh-Gal4* and α1 RNAi transgenes, precluding assessment of whether VNC-specific α1 depletion alters sleep. Other RNAi transgenes exhibited similar lethality when combined with *tsh-Gal4*, suggesting synthetic lethality in certain *tsh-Gal4* heterozygous backgrounds; *tsh-Gal4* is homozygous lethal. In contrast, expression of β1 RNAi using *tsh-Gal4* did not alter viability and significantly decreased total sleep duration with respect to controls bearing only Gal4 or RNAi transgenes (Figures 7B and 7C). Sleep bout number was increased while bout length was reduced (Figures S7A and S7B), indicating that depleting β1 in *tsh-Gal4*- expressing neurons reduces the duration and maintenance of sleep. Together, these data strongly suggest that β1 is required within the VNC for normal sleep.

**Figure 7:**
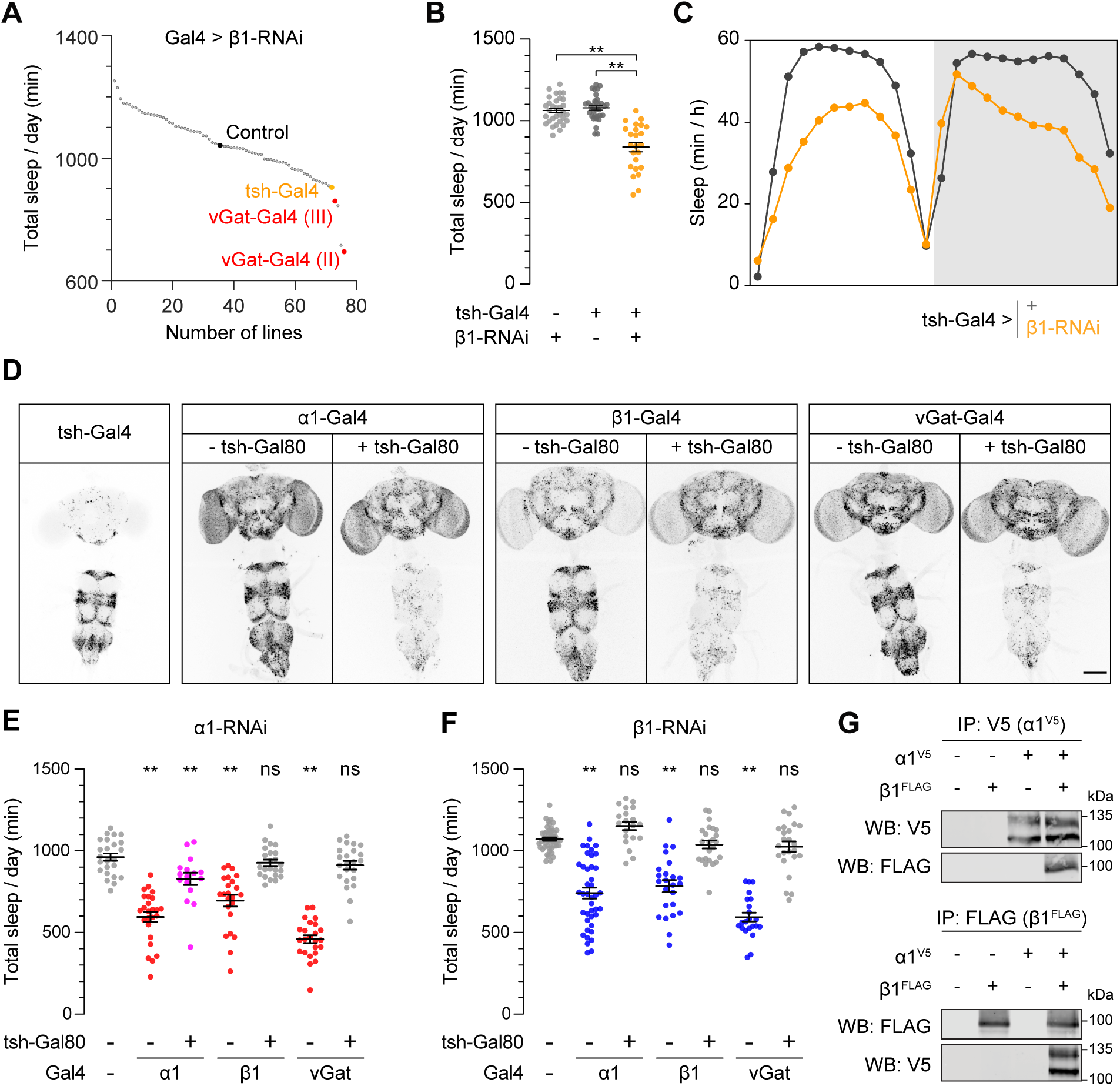
α1 and β1 regulate sleep through ventral nerve cord neurons. **(A)** Total sleep for Gal4 lines expressing β1-RNAi. Black dot, *UAS-β1-RNAi; UAS-dcr2* control animals lacking Gal4. *vGat-Gal4 (II)* and *vGat-Gal4 (III)* are 2nd and 3rd chromosome insertions respectively. **(B)** Depletion of β1 using *tsh-Gal4* elicits short sleep. n= 24-32. **(C)** Average sleep profiles of animals shown in **(B)**. **(D)** *tsh-Gal4*-driven nuclear *UAS-RedStinger* signal is largely restricted to the ventral nerve cord (VNC). *tsh-Gal80* suppresses VNC expression of *UAS- RedStinger* driven by *α1^Gal4^*, *β1^Gal4^*, and *vGat-Gal4*. Scale bar represents 100 µm. **(E)** *tsh-Gal80* restores normal sleep to animals expressing α1 RNAi under the control of *α1^Gal4^*, *β1^Gal4^*, and *vGat- Gal4*. n = 16-25. **(F)** *tsh-Gal80* restores normal sleep to animals expressing β1 RNAi under the control of *α1^Gal4^*, *β1^Gal4^*, and *vGat-Gal4*. n = 20-47. **(G)** α1 and β1 assemble into a heteromeric nAChR in the VNC *in vivo*. Reciprocal co-immunoprecipitations (IP) and Western blots (WB) of α1 and β1 from thorax lysates prepared from *β1^FLAG^ α1^V5^* heterozygotes or control animals. **(B, E and F)** Bars indicate mean ± SEM. One-way ANOVA and Tukey post hoc tests; **p < 0.01; ns, not significant.

To further investigate sleep-regulatory functions of α1 and β1 that arise in the VNC and to determine whether α1 and β1 might impact sleep through GABAergic VNC neurons, we used combinatorial genetic manipulations involving Gal4 and *tsh-Gal80*, a transgene expressing the Gal80 suppressor specifically in the VNC. We first determined whether *tsh-Gal80* could efficiently suppress the expression of UAS transgenes in VNC neurons, by expressing nuclear RedStinger^81^ using *α1^Gal4^*, *β1^Gal4^*, and *vGat-Gal4* in the presence or absence of *tsh-Gal80*. We observed that *tsh-Gal80* robustly suppressed RedStinger signal specifically in VNC neurons for each of these Gal4 drivers, while sparing expression in the brain (Figure 7D). Next, we measured the effects on sleep when the same Gal4 drivers were used to express RNAi against α1 or β1 in the presence or absence of *tsh-Gal80*. Strikingly, the short-sleep phenotypes caused by α1 and β1 depletion using *α1^Gal4^*, *β1^Gal4^*, and *vGat-Gal4* were strongly or fully suppressed by *tsh-Gal80* (Figures 7E and 7F). These results indicate that α1 and β1 depletion solely within the brain—including within GABAergic neurons—is insufficient to cause short sleep phenotypes. Instead, these results support a critical sleep-promoting function for both α1 and β1 that originates in GABAergic VNC neurons.

Finally, to assess whether α1 and β1 assemble into a heteromeric nAChR in the VNC, as in the brain (Figure 4A), we prepared lysates from isolated thorax preparations that contain the VNC and performed reciprocal co-immunoprecipitations. In animals bearing both α1^V5^ and β1^FLAG^, we observed that α1^V5^ co-immunoprecipitated with anti-FLAG, and conversely, that β1^FLAG^ co- immunoprecipitated with anti-V5 antibody (Figure 7G), indicating that α1 and β1 form a heteromeric nAChR within the VNC. The assembly of α1 and β1 in VNC neurons, combinatorial genetic manipulations of these subunits in the VNC (Figures 7D–7F), and pharmacological rescue of their sleep phenotypes with a GABA agonist (Figure 6), strongly support a vital sleep-promoting function for GABAergic VNC neurons that express heteromeric α1/β1 receptors.

## DISCUSSION

In this study, we identify a discrete heteromeric nicotinic acetylcholine receptor that impacts sleep through GABAergic neurons of the VNC, a structure critical for motor and sensory function. We hypothesize that cholinergic excitation of α1^+^/β1^+^ GABAergic VNC neurons evokes GABA release within the VNC and promotes sleep by inhibiting motor neurons, sensory neurotransmission, or both, serving as a vital relay from sleep regulatory circuits in the brain (Figure 8).

**Figure 8:**
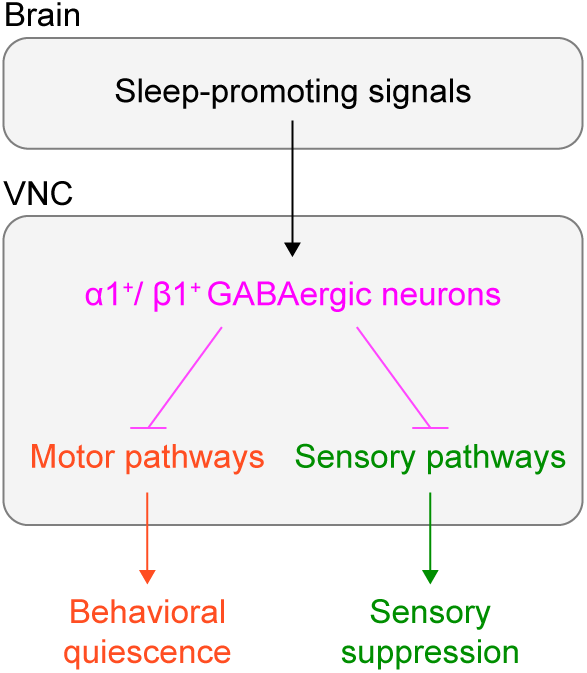
Model for sleep regulation by α1^+^/β1^+^ GABAergic VNC neurons. α1^+^/β1^+^ GABAergic VNC neurons relay cholinergic sleep-promoting signals from the brain to inhibit targets within the VNC including motor and/or sensory pathways, eliciting behavioral quiescence and/or the suppression of sensory stimuli to promote and maintain sleep.

### Motor and sensory systems as loci of sleep regulation and dysfunction

Our findings indicate that heteromeric α1^+^/β1^+^ nAChRs and their expression in GABAergic VNC neurons are essential for sleep regulation. The VNC, homologous to the vertebrate spinal cord, is the ultimate effector of behavioral activity and contains motor neurons, whose activity must be suppressed during sleep. One simple and attractive hypothesis consistent with our findings is that sleep-regulatory GABAergic α1^+^/β1^+^ VNC neurons receive excitatory input from descending cholinergic projections and release GABA to suppress the activity of motor neurons and thereby restrict behavioral activity during sleep (Figure 8), through direct inhibition of motor neurons, indirectly through suppression of excitatory premotor neurons, or through both mechanisms. An analogous mechanism in mammals regulates muscle atonia during REM sleep: glutamatergic neurons in the sublaterodorsal tegmental nucleus (SLD) provide excitatory input to inhibitory interneurons in both the spinal cord and ventral medulla, which in turn silence motor neurons.^82–84^ Anatomical lesions of the SLD, genetic ablation of glutamate release from SLD neurons, or blocking inhibitory GABAergic/glycinergic neurotransmission in spinal interneurons yield similar phenotypes in rodent models: loss of REM atonia, ectopic limb movements, and locomotor activity during sleep.^85,86^ Decreased excitatory input to GABAergic VNC neurons caused by loss of α1 or β1 may cause an analogous disinhibition of motor neurons and ectopic behavioral activity normally associated with wakefulness.

The hypothesis that motor neurons are targets of GABAergic α1^+^/β1^+^ VNC neurons is consistent with the structure and organization of the VNC, as elucidated by recent analysis of its connectome.^87–91^ The vast majority (>80%) of VNC neurons are intrinsic neurons, which have their projections wholly within the VNC and contribute prominently to premotor circuits. Intrinsic neurons provide most of the synaptic inputs to the ∼700 glutamatergic motor neurons, including ∼75% of inputs for leg and wing motor neurons.^88^ The majority of the descending neurons that project from the brain into the VNC are cholinergic, and most of their synaptic outputs are to VNC intrinsic neurons; outputs directly to motor neurons are infrequent.^90,92^

A second, non-mutually exclusive possibility is that GABA released by α1^+^/β1^+^ VNC neurons inhibits sensory afferents that terminate within the VNC or that ascend through the VNC to the brain, suppressing sensory neurotransmission to promote and maintain sleep (Figure 8). Roughly 6,000 sensory neurons have cell bodies in the periphery and project afferents that terminate in the VNC, and another ∼600 sensory afferents ascend through the VNC and terminate in the brain.^87,93^ While analysis of the sensory neuron connectome is ongoing, recent studies have elucidated intrinsic VNC GABAergic neuron subtypes that receive descending input from the brain and provide presynaptic inhibitory input to sensory axons in the context of proprioception and movement.^94^ Whether these or other inhibitory motifs within the VNC might be recruited during sleep to suppress sensory inputs is unknown. In vertebrates, the thalamus is thought to play a primary role in sensory gating during sleep,^95,96^ and analogous mechanisms in the invertebrate brain likely contribute to the suppression of sensory stimuli.^97^ Yet recent evidence suggests that sensory neurons may also be regulated by sleep and contribute in turn to its regulation.^8,9,98^ Sensory afferents that transit the VNC and convey somatosensory stimuli^93^ might be modulated by inhibitory input from GABAergic VNC neurons to promote and maintain the sleep state.

A third non-mutually exclusive possibility is that GABAergic α1^+^/β1^+^ VNC neurons impact sleep by modulating the activity of ascending neurons, which convey information to the brain about behavioral state, including locomotion and resting.^99^ Ascending VNC neurons are an order of magnitude less abundant than intrinsic VNC neurons, but innervate brain regions including those involved in sensory integration and action selection.^99^ If GABAergic α1^+^/β1^+^ VNC neurons provide modulatory input to ascending neurons, reduced α1 or β1 activity could alter signaling to the brain and thereby alter sleep regulation. While GABAergic α1^+^/β1^+^ VNC neurons might themselves be ascending neurons, this seems less likely, given that most GABAergic neurons are intrinsic neurons whose projections reside within the VNC, as noted below.

Identifying the targets of GABAergic α1^+^/β1^+^ VNC neurons in the context of sleep requires functional mapping within GABAergic VNC neurons. Of the 34 neuronal hemilineages that give rise to the adult VNC, 12 are GABAergic and provide significant innervation to leg and wing neuropils.^91,100^ The small number of descending neurons relative to the number of intrinsic VNC neurons,^92^ together with the possibly broad distribution of α1^+^/β1^+^ VNC neurons, suggests that descending sleep-regulatory signals might be amplified within the VNC to broadly inhibit targets and facilitate binary transitions between sleep and waking states. Certain subtypes of GABAergic intrinsic VNC neurons project to all leg connectomes, suggesting they may provide coordinated inhibition of premotor circuits.^90^ Along with functionally defining subpopulations of GABAergic α1^+^/β1^+^ VNC neurons that impact sleep, assessing their activity during sleep-wake cycles and defining their inputs are similarly essential to understanding their impact on behavior.

### A sign-switch mechanism couples excitatory cholinergic input and GABA release and promotes sleep

Although depletion of nAChRs containing α1 and β1, widely distributed excitatory neurotransmitter receptors, might be expected to increase sleep by broadly decreasing neuronal excitation, α1 or β1 depletion strongly curtails sleep. Our findings suggest a “sign-switch” mechanism as the resolution to this paradox: α1 and β1 promote sleep by coupling excitatory cholinergic input to inhibitory GABAergic output within the VNC. While one caveat is that *vGat- Gal4* is expressed in additional, non-GABAergic neurons,^78,101^ low concentrations of gaboxadol restore sleep to short-sleeping animals expressing α1 or β1 RNAi against driven by *vGat-Gal4*, supporting a GABAergic mechanism. The administration of gaboxadol to adult animals in these experiments furthermore suggests that α1 and β1 regulate sleep through GABAergic neurons in an ongoing manner.

While our findings favor a sign-switch model, they do not exclude possible contributions from additional mechanisms, including homeostatic upregulation of cholinergic transmission or neuronal activity arising from depletion of α1 and β1. Pharmacological inhibition of nAChRs or acute genetic inhibition of acetylcholine synthesis has been shown to cause transcriptional upregulation of the α7 subunit.^102,103^ However, α1 depletion does not elicit transcriptional upregulation of β1 and vice versa, suggesting that any compensatory mechanisms relevant to sleep would have to involve post-transcriptional mechanisms, other nAChR subunits, or changes at the level of neurons or circuits. The simplest hypothesis consistent with our findings is that short sleep phenotypes caused by α1 or β1 depletion result from reduced cholinergic excitatory input to GABAergic neurons whose activity promotes sleep.

### Functional dissection of heteromeric nAChRs in vivo

Elucidating functions of heteromeric neurotransmitter receptors within the nervous system is an enduring challenge due to the difficulty of manipulating discrete receptors in vivo and linking them to behavior. The in vivo composition, distribution, and function of heteromeric nAChRs in *Drosophila* have remained largely unknown since the cloning of nAChR subunits decades ago.^26–35^ Although the inability to express native fly nAChRs in heterologous systems was recently overcome with expression of a key co-factor,^104^ a second obstacle has been the lack of genetic and biochemical tools for analyzing heteromeric nAChRs in the context of the brain and behavior. The strategies we have used to manipulate α1 and β1 bypass limitations of earlier reagents and enable the function of heteromeric nAChRs to be dissected in vivo. One prong of this strategy is endogenous epitope tagging of multiple receptor subunits, permitting assessment of their physical associations and distribution. Notably, smGFP tags bearing multiple FLAG and V5 epitopes revealed a biochemical association between α1 and β1 which was undetectable with antibodies against endogenous proteins.^59^ Similar endogenous tagging strategies for nAChRs were recently reported in independent studies.^105^ A second prong of this strategy is reciprocal genetic manipulations of other nAChR subunits, enabled by inserting Gal4 into receptor subunit loci. Given that the large size of many receptor loci (e.g. >60 kb for *α1*) limits the ability of transgenes bearing isolated regulatory elements to recapitulate expression patterns, Gal4 insertion alleles may be broadly required to effectively manipulate nAChRs and other heteromeric receptors.

Our findings and recent studies suggest the intriguing possibility that distinct heteromeric nAChRs may assemble in different sleep-regulatory circuits. α2 and β2 were recently shown to be required in octopaminergic neurons to promote sleep in *Drosophila*,^36^ suggesting, together with earlier observations that α2 and β2 associate in head lysates,^59^ that these subunits might assemble into a heteromeric receptor in octopaminergic neurons. An additional interesting possibility is that heteromeric α1/β1 or α2/β2 nAChRs might contain other subunits linked to sleep.^38,106^ The combination of heterologous expression systems, reciprocal manipulations in vivo, and additional intersectional genetic strategies may allow *Drosophila* to serve as a powerful system for analyzing nAChRs that impact sleep and other behaviors. Applying analogous strategies in rodent models for nAChR mutations linked to sleep-related epilepsy^107,109,111^ might similarly elucidate heteromeric nAChRs relevant for pathological sleep disruptions in humans. Such findings could have therapeutic relevance, as they might suggest specific pharmacological targets for the treatment of sleep disorders.

## Methods

### Fly lines and rearing conditions

Flies were reared on standard fly food at 25**°**C in alternating 12-hour light-dark (LD) cycles. The following stocks were obtained from the Bloomington Drosophila Stock Center: *w*^1118^ (RRID: BDSC_5905), *elav^C^*^155^–*Gal4; UAS-dcr2* (RRID: BDSC_25750)*, y*^1^ *w*^67c23^*; sna^Sco^/CyO, P{Crew}DH1* (RRID: BDSC_1092), *α1-MiMIC*^453^ (RRID: BDSC_42295), *vGat-Gal4 (III)* (RRID: BDSC_58409), *tsh-Gal4* (RRID: BDSC_3040), Gal4 lines bearing putative *α1* regulatory fragments (R22A03, RRID: BDSC_48962; R22G02, RRID: BDSC_48997; R22G10, RRID: BDSC_49000; 22E11, N/A; R22A09, N/A; R23B01, N/A)*. UAS-α1-RNAi* (#48159 and #48162) and *UAS-β1-RNAi* (#106570) were obtained from the Vienna Drosophila Resource Center. UAS- α1-RNAi (#5610R-3) and UAS-α4-RNAi (#12414R-3) were obtained from the National Institute of Genetics Fly Stock Center. *tsh-Gal80* and *UAS-myrGFP-2A-RedStinger* were gifts from Treisman lab and Ganetzky lab, respectively.

### Molecular Biology

To generate *α1^attP^* and *β1^attP^*, the first coding exons of α1 and β1 were excised using flanking sgRNAs and the CRISPR/Cas9 system (Figure S2A). To generate plasmids expressing α1 and β1 sgRNAs, oligonucleotides were annealed and cloned into BbsI-digested pU6-BbsI-chiRNA (Addgene, RRID: Addgene_45946).^108^ Plasmids for homology-directed repair (pHD-DsRed-α1^attP^ and pHD-DsRed-β1^attP^) were generated by amplifying homology arms from genomic DNA extracted from the same strain used for targeting and cloned into pHD-DsRed-attP^58^ (Addgene, RRID: Addgene_51019). Cocktails comprising two sgRNA plasmids and the HDR plasmid were injected into *y*^2^ *cho*^2^ *v*^1^*; Sp/CyO, P{nos-Cas9, y^+^, v^+^}2* embryos^110^ by BestGene Inc. Animals bearing correct CRISPR targeting events at the α1 and β1 loci were isolated by selecting for eye-specific DsRed expression and subsequently validated by PCR. The loxP-flanked 3*×*P3-DsRed marker was subsequently excised by crossing with animals expressing Cre (*y*^1^ *w*^67c23^*; snaSco/CyO, P{Crew}DH1*, RRID: BDSC_1092); precise excision was confirmed by PCR. *α1^attP^* deletes 1322 bp (R6: 3R 24420839–24422160), and *β1^attP^* deletes 1931 bp (R6: 3L 4428458– 4430388).

To generate smGFP-tagged alleles of α1 and β1 (*α1^V5^* and *β1^FLAG^*), exons encoding the large intracellular loop between the third and fourth transmembrane domains were modified by HDR (Figure S3). Insertion sites were chosen based on the absence of sequence conservation with other insect α1 or β1 subunits and the lack of potential post-translational modification sites.^112,113^ smGFP tags were flanked by GGGGS linker sequences and inserted after residues A418 and G424 for α1 and β1, respectively.

*α1^Gal4^* was generated from *α1-MiMIC*^453^ (RRID: BDSC_42295) using the MiMIC/Trojan system.^76^ Insertion of the T2A-Gal4-polyA Trojan cassette into the third intron of α1 yields bicistronic expression of Gal4 and a truncated α1 protein predicted to lack all transmembrane domains and to be nonfunctional.

To generate *β1^Gal4^*, a plasmid bearing an *attB* site, a 3*×*P3-GFP selection marker, and a modified β1 exon with insertions of Gal4 and an SV40 polyadenylation signal immediately downstream of the β1 start codon (pGFP-attB-β1^Gal^^4^) was injected into *β1^attP^* embryos expressing germline-specific phiC31 recombinase (BestGene Inc). Animals with plasmid integration at the attP site of *β1^attP^* (*β1^Gal4+GFP^*) were selected by eye-specific 3*×*P3-GFP expression and validated by PCR. The loxP-flanked 3*×*P3-GFP marker and plasmid backbone were subsequently excised with Cre recombinase (*y*^1^ *w*^67c23^*; snaSco/CyO, P{Crew}DH1*, RRID: BDSC_1092). Excision of the 3*×*P3-GFP marker was confirmed by PCR. *β1^Gal4^* expresses Gal4 instead of β1 protein, yielding a presumed β1 null allele.

To generate *UAS-α1* and *UAS-β1*, RNA was extracted from head lysates of *w*^1118^ (RRID: BDSC_5905) animals using TRIzol (Thermo Fisher Scientific) and purified using RNeasy mini spin columns (Qiagen). cDNA was synthesized with d(T)23VN primer using SuperScript II Reverse Transcriptase (Thermo Fisher Scientific). α1 and β1 coding sequences corresponding respectively to α1-PA and β1-PA were amplified with primers containing EcoRI and XhoI sites and cloned into EcoRI-XhoI digested pUASTattB.^114^ pUASTattB-α1 and pUASTattB-β1 were integrated into *attP40* (BestGene Inc).

### Sleep and circadian assays

One- to four-day-old flies eclosing from cultures entrained in LD cycles were loaded into glass tubes (Trikinetics) and assayed for 5–7 days at 25**°**C in LD cycles using DAM2 monitors (Trikinetics). Male flies were assayed on standard fly food containing cornmeal, agar, and molasses. For pharmacological manipulation of sleep, female flies were raised on standard fly food and assayed on food containing 5% sucrose, 2% agar, and either water vehicle or gaboxadol (Sigma-Aldrich) at 0.005 mg/mL or 0.01 mg/mL. The first 36–48 hours of data were discarded to permit acclimation and recovery from CO2 anesthesia, and four to five integral days of data were analyzed using custom Matlab software.^68^ Locomotor data was collected in one-minute bins, and periods of five minutes or more of inactivity were defined as sleep.^115,116^ Dead animals were excluded from analysis by a combination of automated filtering and visual inspection of locomotor traces.

To assess circadian rhythmicity, animals were entrained to LD cycles and subsequently released into constant darkness (DD) for five days. Behavioral rhythmicity was analyzed using *χ*^2^ periodogram analysis in ClockLab (ACTIMETRICS, RRID: SCR_014309). For sleep deprivation experiments, animals were loaded in DAM2 monitors and placed in an apparatus for programmed mechanical stimulation (VMP, TriKinetics). After 48 h acclimation, animals were allowed to sleep undisturbed for one day (baseline, B) prior to sleep deprivation on the second day during the second half of the night (ZT18–ZT24) using one second of gentle mechanical vibration every minute at a maximum speed setting of 3 on the VMP apparatus (sleep deprivation, SD). Animals were allowed to sleep unperturbed on the third day (recovery, R). Animals which lost 70% of sleep or more were used to calculate the amount of sleep lost and sleep rebound as follows: Sleep lost = Sleep (ZT18–ZT24, B) - Sleep (ZT18–ZT24, SD) Sleep regained = Sleep (ZT0–ZT6, R) -Sleep (ZT0–ZT6, B)

### Quantification of α1 and β1 mRNA levels with qRT-PCR

RNA extraction and cDNA synthesis were performed as described above. Transcript levels of α1, β1, and RPS3 were quantified in triplicate from cDNA using SsoAdvanced Universal SYBR Green Supermix (Bio-Rad). α1 and β1 transcript levels were normalized using the average Ct value of RPS3 for each experiment. Two independent experiments were performed. qRT-PCR was performed with the following primers:

α1 forward: 5’-CGCCGATCTCAGTCCTACGTTCGAGA-3’

α1 reverse: 5’-GCAAGCGATCGCGAAGATCCACAGA-3’

β1 forward: 5’-GGTGCGTGACCTCTTTCGAGGCTA-3’

β1 reverse: 5’-CCACTGCAGCTGGTAGTCGTACCAA-3’

RPS3 forward: 5’-CGAACCTTCCGATTTCCAAGAAACGC-3’

RPS3 reverse: 5’-ACGACGGACGGCCAGTCCTCC-3’

### Biochemistry

Protein extracts were prepared from frozen sieved heads or frozen isolated thoraxes by manual pestle homogenization in ice-cold NP-40 lysis buffer (50 mM Tris pH 7.6, 150 mM NaCl, 0.5 % NP-40) supplemented with protease inhibitors (Sigma-Aldrich). To isolate thoraxes, bodies separated by sieving were placed on an aluminum block cooled by dry ice and a dry-ice cooled razor blade was used to remove and discard heads and abdomens from individual bodies.

Lysates were centrifuged at 4°C at 15,000 g for 15 minutes and supernatants were extracted. Total protein levels in each sample were quantitated in duplicate (DC protein assay, Bio-Rad). For head samples, 1.5 mg of protein in a total volume of 100 microliters was incubated at 4°C overnight with 20 microliters of mouse anti-V5 (1:100, Novus, RRID: AB_1625981) or rat anti-FLAG (1:100, Sigma-Aldrich, RRID: AB_261888); thorax samples were scaled identically, using 6 mg of protein in a total volume of 200 microliters and 40 microliters of antibody. Head and thorax samples were respectively immunoprecipitated by incubation with 20 or 40 microliters of a 50% slurry of protein G-Sepharose 4B (Thermo Fisher Scientific) at 4°C for 4 hours followed by four 15 min washes with ice-cold NP-40 lysis buffer. Immunoprecipitated samples were resolved by Tris-SDS-PAGE and transferred to nitrocellulose membranes (Santa Cruz Biotech). Membranes were blocked in Odyssey Blocking Buffer (LI-COR) for one hour at room temperature and subsequently incubated overnight at 4**°**C in a cocktail containing Odyssey Blocking Buffer, 0.1 % Tween 20, and mouse anti-V5 (1:2000) or rat anti-FLAG (1:500). After four 5 min washes in TBST (150 mM NaCl, 10 mM Tris pH 7.6, and 0.1 % Tween20), membranes were incubated in the dark at room temperature for one hour in a cocktail containing Odyssey Blocking Buffer, 0.1 % Tween 20, 0.01 % SDS, and the following secondary antibodies: donkey anti-rat Alexa 680 (1:30,000, Thermo Fisher Scientific, RRID: AB_2534014) and donkey anti-mouse Alexa 790 (1:30,000, Jackson Immunoresearch, RRID: AB_2340701). After four 5 min washes in TBST and a wash in TBS (150 mM NaCl, 10 mM Tris pH 7.6) for 5 min, membranes were scanned with an Odyssey CLx imager (LI-COR, RRID: SCR_014579).

### Immunohistochemistry

For immunohistochemistry of adult fly brains with attached ventral nerve cords, whole animals were rinsed with PBS (137 mM NaCl, 2.7 mM KCl, 10 mM Na2HPO4, 1.8 mM KH2PO4) containing 0.2 % Triton X-100 (PBST), dissected in ice-cold PBS, and fixed either at room temperature for 30–60 min with 4% paraformaldehyde in PBS or overnight at 4°C with 2% paraformaldehyde in PBS. After three brief rinses with PBST, samples were washed three times in PBST for 15 min at room temperature. Samples were blocked for 1h at room temperature in PBST containing 5% normal donkey serum (Lampire Biolgical) and subsequently incubated for 2–3 days at 4°C in primary antibody cocktail diluted in blocking solution. Following three brief rinses and three 20 min washes with PBST at room temperature, samples were incubated in secondary antibody cocktail in blocking solution for 1–2 days at 4°C. Samples were then washed three times with PBST at room temperature and mounted onto microscopy slides with Vectashield (Vector Laboratories, RRID: AB_2336789). Primary antibodies were mouse anti-V5 (1:400), rat anti-FLAG (1:200), rabbit anti-PER (1:50, Sehgal lab), mouse anti-PDF (1:20, DSHB, RRID: AB_760350), chicken anti-GFP (1:500, Aves Labs, RRID: AB_10000240), rabbit anti-GFP (1:3,000, Life Technologies, RRID: AB_221569), and rabbit anti-DsRed (1:1,000; Takara, RRID: AB_10013483). Secondary antibodies, all used at 1:1,000, were donkey anti-chicken Alexa 488 (Jackson Immunoresearch, RRID: AB_2340375), donkey anti-rabbit Alexa 488 (Life Technologies, RRID: AB_2535792), donkey anti-rat Alexa 488 (Life Technologies, RRID: AB_2535794), donkey anti-mouse Alexa 568 (Life Technologies, RRID: AB_2534013), and donkey anti-rabbit Alexa 568 (Life Technologies, RRID: AB_2534017). For staining of smGFP- tagged animals, samples were incubated in 0.8 μg/mL DAPI in PBST for 30 min prior to mounting. For staining of circadian pacemaker neurons, animals entrained to LD cycles were released into constant darkness and collected on the second day of constant darkness at CT0, CT6, CT12, and CT18 for dissection, fixation, and staining as described above.

All imaging was performed on Zeiss LSM510 or LSM800 confocal microscopes. For Figures 2B and 3C, confocal slices were captured using 40*×* objective at 1024 x 1024 resolution. Images in Figure 3B were captured using 20*×* objective at 512*×*512 resolution. Images in Figure 7D were captured using a 10*×* objective at 1024 x 1024 resolution. Z-stack maximal projections were generated using Fiji software (FijiSc, RRID: SCR_002285).

### Quantification and statistical analysis

Statistical comparisons of total sleep, daytime sleep, nighttime sleep, sleep bout number, activity per waking minute, and mRNA abundance were conducted using one-way ANOVA and Tukey or Dunnett’s post-hoc tests. Kruskal-Wallis and Dunn’s post-hoc tests were used for comparisons of sleep bout length. For pharmacological manipulations of sleep, sleep parameters and locomotor activity were compared with two-way ANOVA and Tukey post-hoc tests.

## ACKNOWLEDGEMENTS

We thank E. Gundelfinger and U. Thomas for sharing antibodies against α1 and β1; J. Treisman and B. Ganetzky for sharing *tsh-Gal80* and *UAS-myrGFP-2A-RedStinger*, respectively; and the Bloomington Drosophila Stock Center, the Vienna Drosophila Resource Center, and the Fly Stock center of the National Institute of Genetics for fly lines. We thank S. Burden, C. Desplan, N. Ringstad, and members of the Stavropoulos lab for comments on the manuscript. This work was supported by grants from the National Institutes of Health (NS112844), the Mathers Foundation, Whitehall Foundation grant 2013-05-78, fellowships from the Alfred P. Sloan and Leon Levy Foundations, a NARSAD Young Investigator Award from the Brain and Behavior Foundation, the J. Christian Gillin, M.D. Research Award from the Sleep Research Society Foundation, and a Career Scientist Award from the Irma T. Hirschl/Weill-Caulier Trust to N.S.

## AUTHOR CONTRIBUTIONS

T.Y., H.A., H.H., I.C., C.L., and N.S. designed experiments, conducted experiments, and analyzed data; T.Y. and N.S. wrote the paper; N.S. supervised the project.

## DECLARATION OF INTERESTS

The authors declare no competing interests.

---

**Figure S1:**
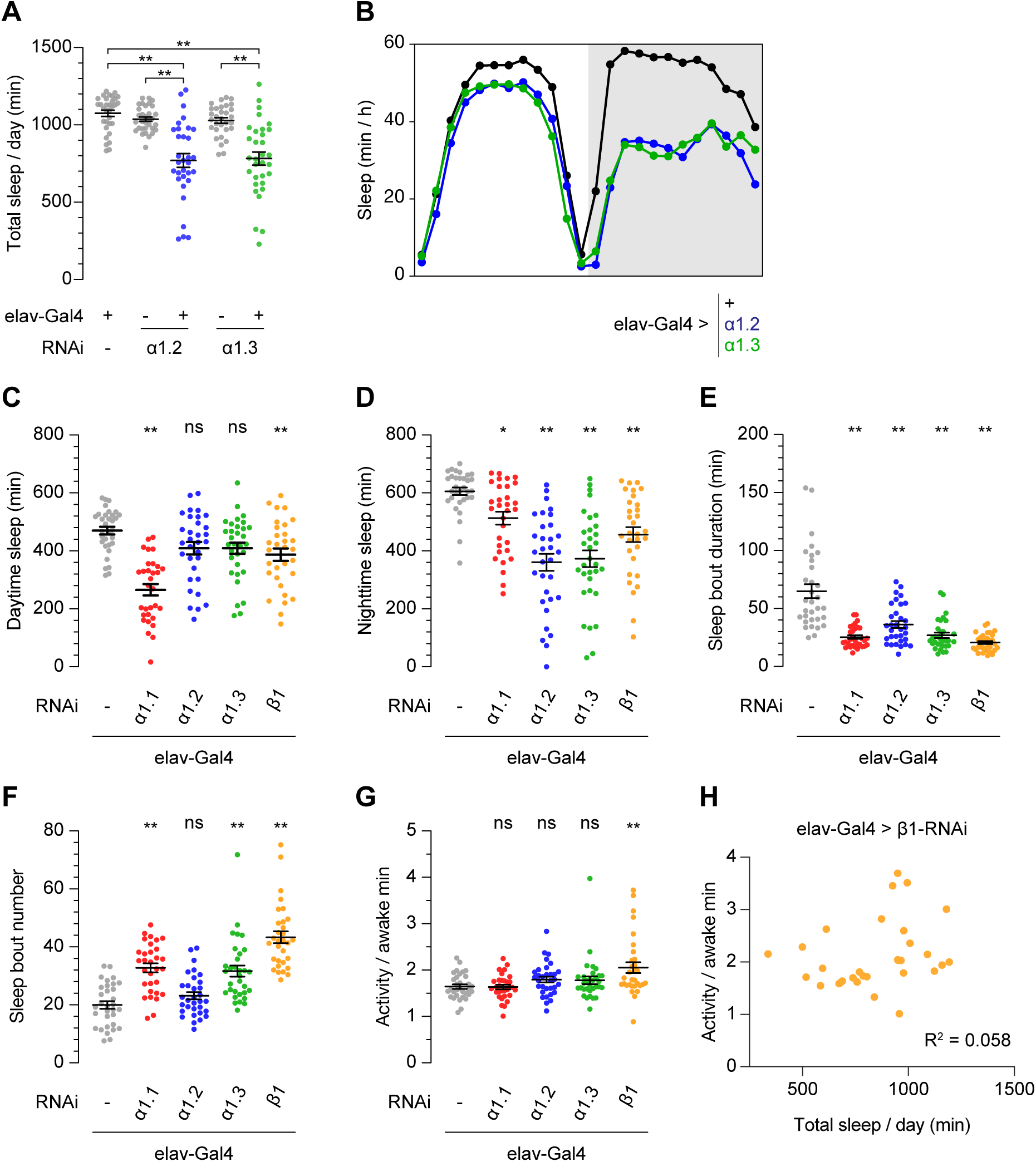
Additional sleep parameters for α1 and β1 RNAi. **(A)** Pan-neuronal depletion of α1 using additional RNAi lines (α1.2, VDRC 48159; α1.3, VDRC 48162) elicits short sleep similar to α1 RNAi (α1.1: Japan 5610R-2) shown in Figures 1B and 1C. n = 31-32. Bars indicate mean ± SEM. One-way ANOVA and Tukey post hoc tests; **p < 0.01. **(B)** Average sleep profiles of animals shown in **(A)**. **(C)** Daytime sleep. **(D)** Nighttime sleep. **(E)** Sleep bout duration. **(F)** Sleep bout number per day. **(G)** Activity per awake minute. n = 31-32. Bars indicate mean ± SEM. Significance is compared to elav-Gal4 control with one-way ANOVA and Dunnett’s tests **(C, D, F, and G)** and Kruskal-Wallis and Dunn’s tests **(E)**; *p < 0.05 and **p < 0.01; ns, not significant. **(H)** Scatterplot of activity and sleep in animals expressing pan-neuronal β1 RNAi.

**Figure S2:**
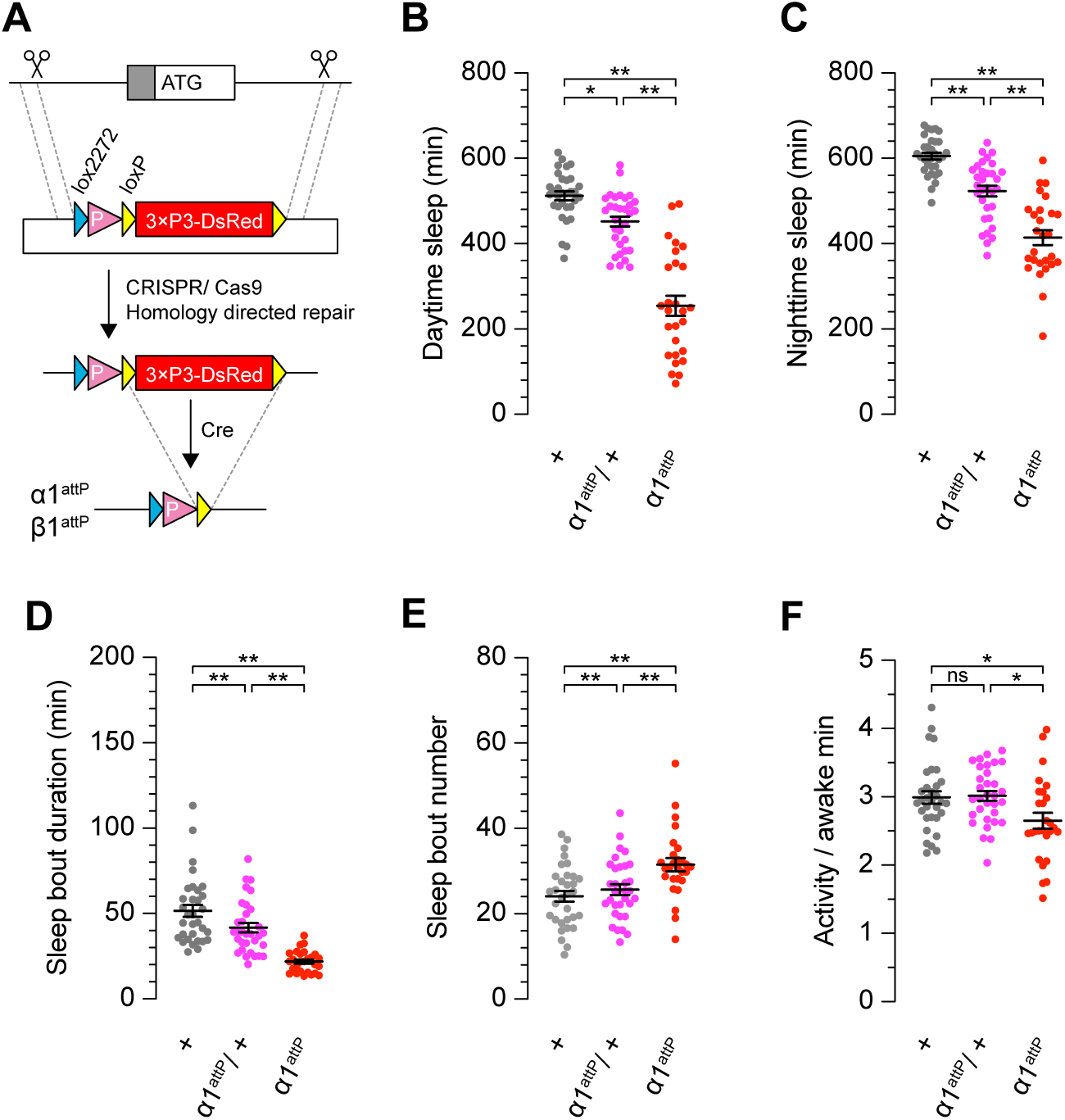
Generation and characterization of *α1^attP^*. **(A)** *α1^attP^* and *β1^attP^* alleles were generated by CRISPR/Cas9 targeting and subsequent excision of the 3×P3-DsRed marker with Cre recombinase. **(B)** Daytime sleep. **(C)** Nighttime sleep. **(D)** Sleep bout duration. **(E)** Sleep bout number per day. **(F)** Activity per awake minute. n = 27-32. Bars indicate mean ± SEM. One-way ANOVA and Tukey post hoc tests **(B, C, E, and F)** or Kruskal-Wallis and Dunn’s tests **(D)**; *p < 0.05; **p < 0.01; ns, not significant.

**Figure S3:**
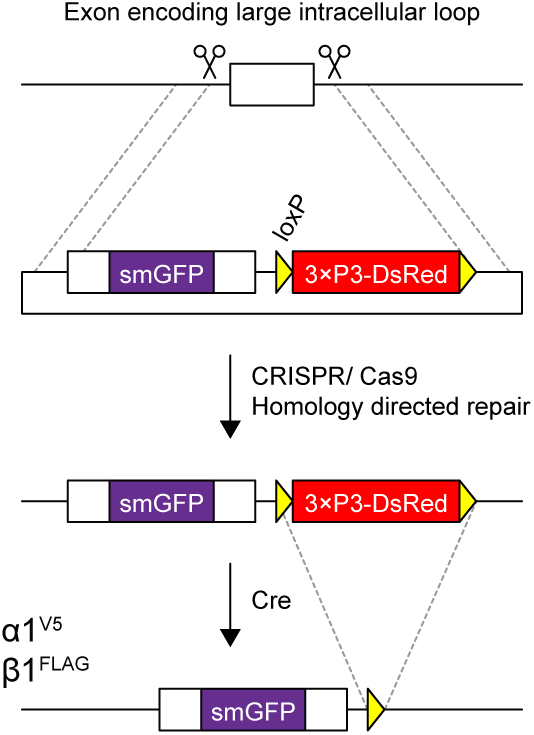
Generation of *α1^V5^* and *β1^FLAG^*. *α1^V5^* and *β1^FLAG^* alleles were generated by CRISPR/Cas9 targeting and subsequent excision of the 3×P3-DsRed marker with Cre recombinase.

**Figure S4:**
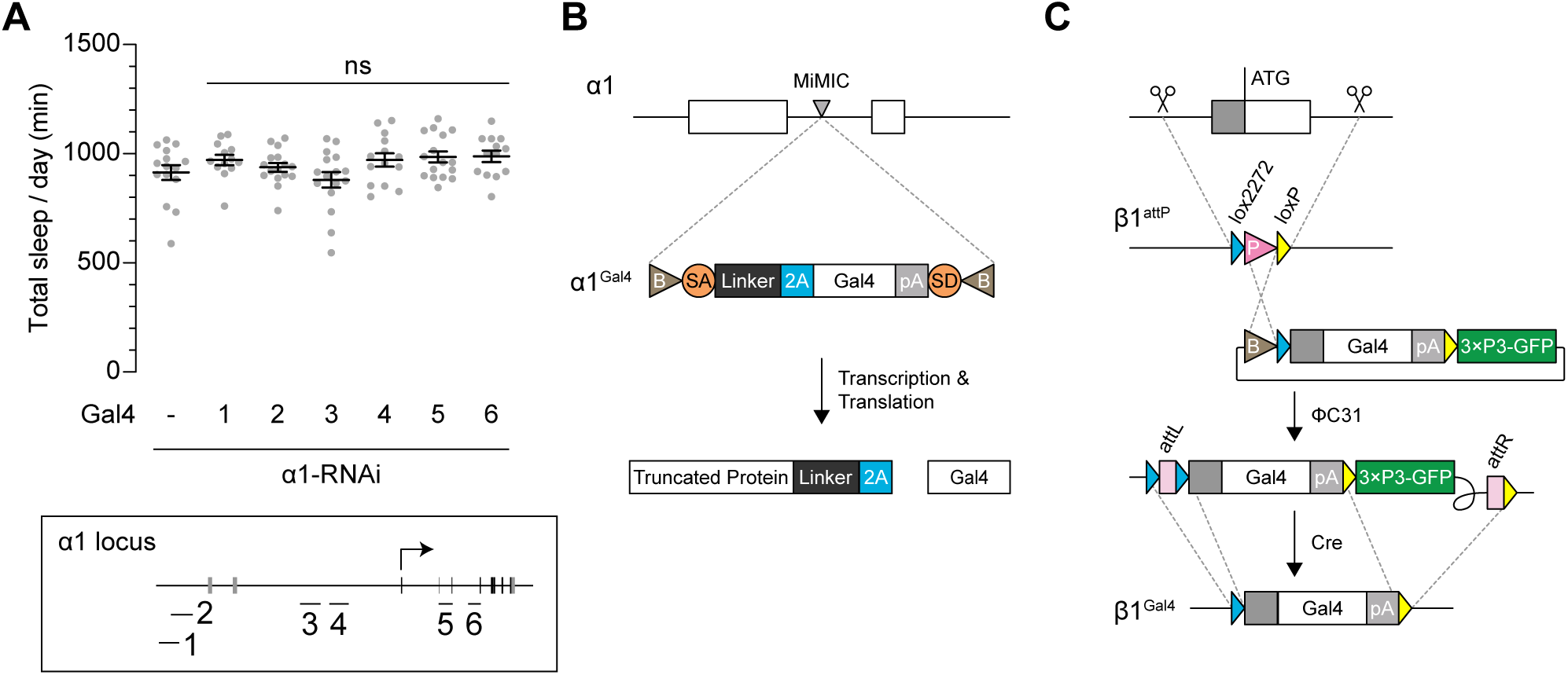
Generation of *α1^Gal4^* and *β1^Gal4^* knock-in Gal4 alleles. **(A)** α1 RNAi expressed with *Gal4* lines bearing putative regulatory fragments from the *α1* locus does not cause short sleep. n = 13-16. Bars indicate mean ± SEM. One-way ANOVA and Dunnett’s tests; ns, not significant with respect to control. Bottom panel indicates locations of *α1* fragments. Gray and black rectangles represent UTR and coding sequence of α1, respectively. **(B)** *α1^Gal4^* was generated by replacing a MiMIC insertion in the *α1* locus with a Trojan Gal4 cassette. The 2A peptide allows bicistronic expression of Gal4 in neurons expressing α1. **(C)** *β1^Gal4^* was generated in two steps from *β1^attP^* (see Figure S2A). In the first step, ΦC31 recombinase was used to integrate a plasmid containing the first coding exon of *β1*, modified such that the *β1* start codon is fused to Gal4. In the second step, the 3×P3-GFP selection marker and vector backbone were removed by Cre recombinase, yielding *β1^Gal4^*. **(B and C)** P, attP; B, attB; SA, splice acceptor; SD, splice donor; 2A, 2A peptide sequence; pA, polyA.

**Figure S5:**
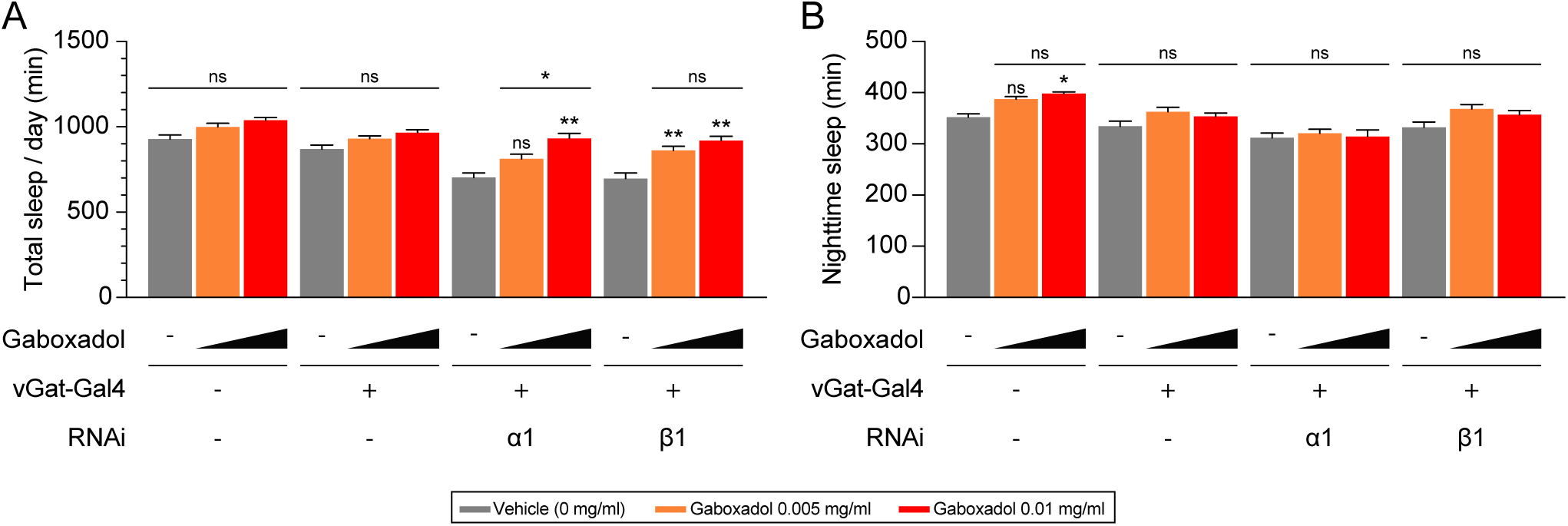
Effects of augmented GABAA receptor activity on nighttime sleep and total sleep. **(A)** Total sleep. **(B)** Nighttime sleep. n = 21-22 as in Figure 6. Mean ± SEM is shown. Two-way ANOVA and Tukey post hoc tests; *p < 0.05; ns, not significant.

**Figure S6:**
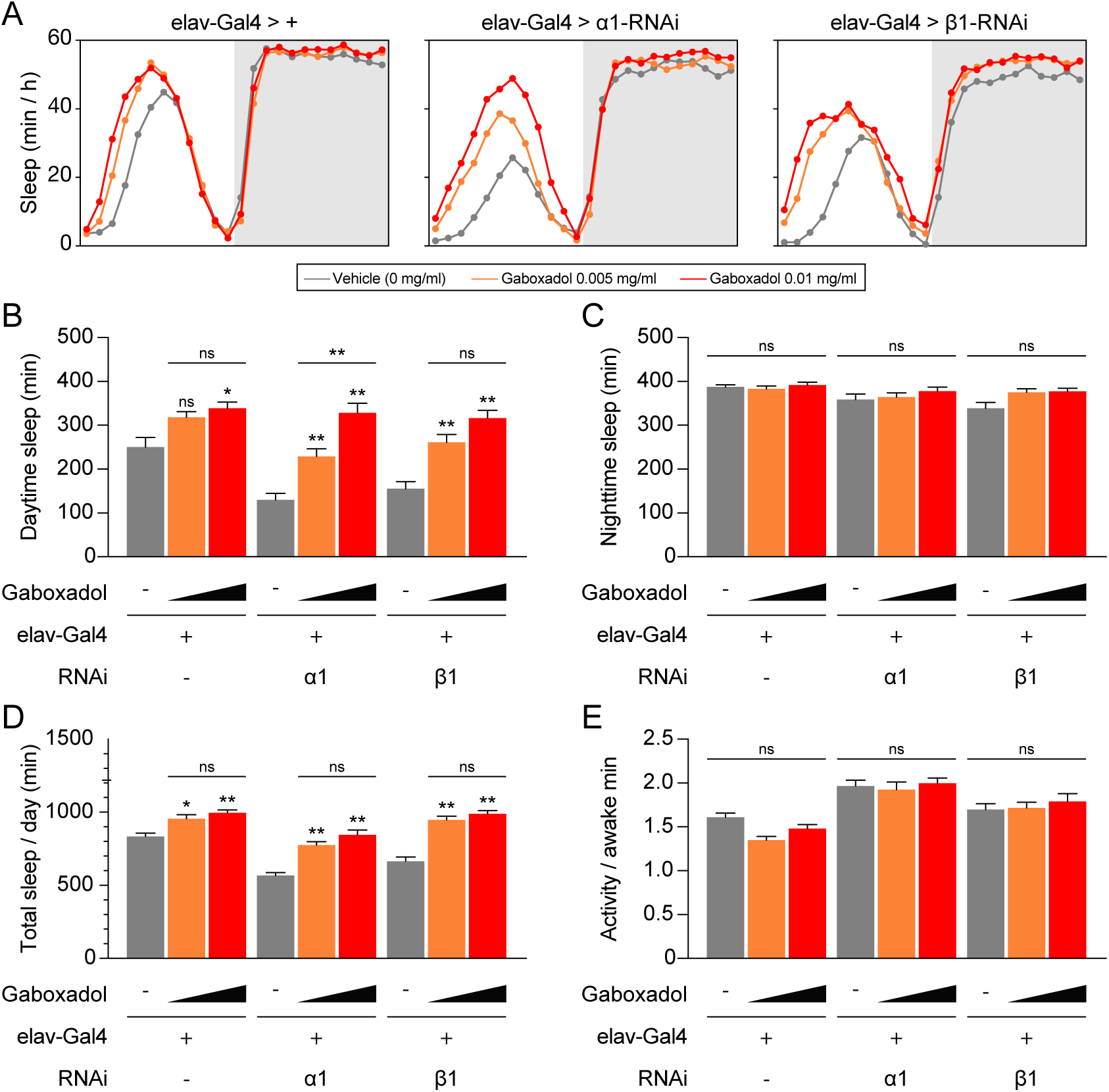
The GABAA receptor agonist gaboxadol complements the sleep deficits of animals expressing pan-neuronal α1 or β1 RNAi. **(A)** Average sleep profiles of indicated genotypes exposed to gaboxadol. **(B)** Daytime sleep. **(C)** Nighttime sleep. **(D)** Total sleep. **(E)** Activity per awake minute. n = 21-22. For **(B–E)**, mean ± SEM is shown. Two-way ANOVA and Tukey post hoc tests; *p < 0.05; **p < 0.01; ns, not significant.

**Figure S7:**
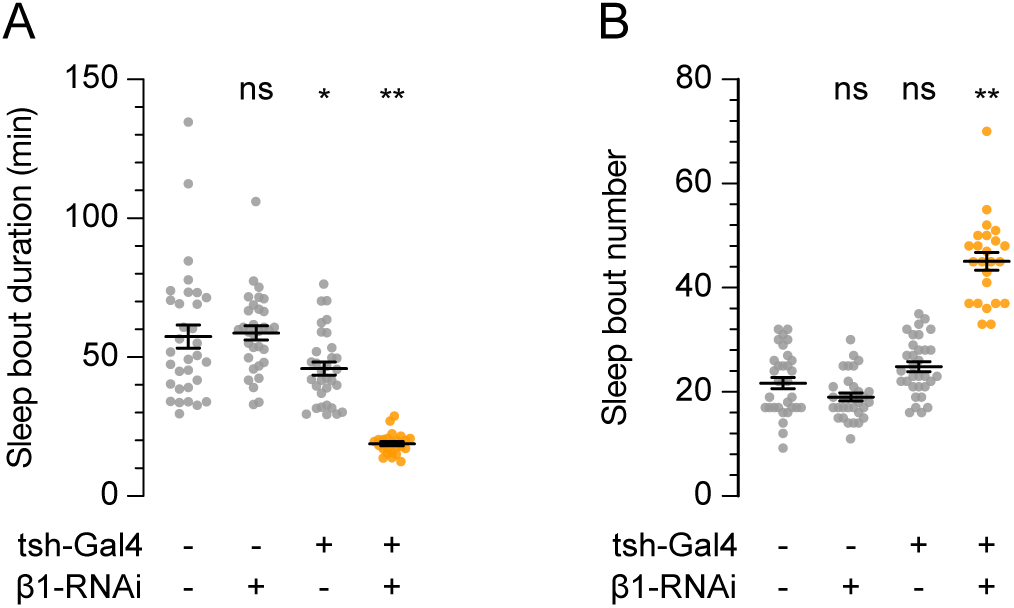
β1 depletion in *tsh-Gal4*-expressing neurons causes fragmented sleep. **(A)** Sleep bout duration. **(B)** Sleep bout number per day. n = 24-32. Bars indicate mean ± SEM. Kruskal-Wallis and Dunn’s tests **(A)** and one-way ANOVA and Tukey post hoc tests **(B)**; *p < 0.05; **p < 0.01; ns, not significant.

## Notes

### Competing Interest Statement

The authors have declared no competing interest.

